# *RHOA* Deletion Downregulates CD19 and Promotes Dysfunctional Immune Microenvironments in CAR-T Resistant B-Cell Lymphoma

**DOI:** 10.1101/2025.02.27.640687

**Authors:** Austin D. Newsam, Bachisio Ziccheddu, Abdessamad A. Youssfi, Venu Venkatarame Gowda Saralamma, Yitzhar E. Goretsky, Isaiah Sheffield-Veney, Paola Manara, Daniel E. Tsai, Adithi Jeevan, Santiago Foos-Russ, David M. Suissa, Marco Vincenzo Russo, Nikolai Fattakhov, David Carmona-Berrio, Natalia Campos Gallego, Caroline A. Coughlin, Anya K. Sondhi, Evan R. Roberts, Jay Y. Spiegel, Juan Pablo Alderuccio, Catalina Amador, Daniel Bilbao, David G. Coffey, Michael D. Jain, Francesco Maura, Frederick L. Locke, Jonathan H. Schatz

## Abstract

CD19-directed chimeric antigen receptor (CAR)-T cells are breakthrough therapies for aggressive B-cell lymphomas, but less than half of patients achieve durable responses. We previously showed through whole-genome sequencing of tumors from CAR-T-treated patients that deletions of *RHOA* (3p21.31) are enriched in cases progressing after treatment. *RHOA*’s roles in resistance and pathogenesis are poorly defined, despite loss-of-function alterations that occur in ∼20% of newly diagnosed large B-cell lymphoma (LBCL) cases. We created RHOA-deficient LBCL systems and confirmed cell-intrinsic loss of response to CAR-19 in vitro and in vivo driven by CD19 downregulation. Impact on CD19, however, was variable and would not explain selection for *RHOA* deletion in newly diagnosed cases. We therefore created RHOA-deficient tumors in immunocompetent mice and found remarkable correlation with dysfunctional lymphoma microenvironment (LME) signatures in CAR-19 resistant patients. These LMEs are marked by a type 1-like immune infiltrate with terminally exhausted CD8 T cells, Th1-like CD4 cytotoxic lymphocytes (CTLs), and increased production of interferon gamma (IFNγ). RHOA-deficient tumor cells themselves have significantly impaired IFNγ responses, providing resistance to CD8 T cell clearance by way of diminished induction of major histocompatibility complex class I (MHC-I). These findings support a model that depletion of healthy effector populations by RHOA-deficient lymphoma is a key driver of immune dysfunction thwarting CAR-19 clinical responses. Overall, we describe for the first time how a single-gene alteration found recurrently in CAR-19-resistant LBCL contributes to treatment failures.

## INTRODUCTION

CD19-targeted autologous chimeric antigen receptor T-cell (CAR-19) immunotherapies are potentially curative for patients with relapsed or refractory (r/r) aggressive B-cell lymphomas.^1–3^ Durable complete response (CR) in >30% represent a major breakthrough, but patients with unsatisfactory responses, unfortunately still the majority, have poor prognosis.^4^ Detailed comprehension of resistance is an urgent priority to inform future immunotherapies. While characteristics of CAR-19 products and lymphoma microenvironments have been well described,^5–8^ the functional roles of lymphoma genomes in thwarting CAR-19 responses are less explored.

We previously showed through whole-genome sequencing (WGS) that genomic factors collectively are stronger predictors of outcome than clinical factors like metabolic tumor volume (MTV) or patient age.^9^ Additional studies implicate, like ours, that genomic complexity and/or alterations to specific genes can predict CAR-19 outcomes.^10–12^ Mechanistic studies of tumor-intrinsic resistance include EZH2 gain-of-function mutations and FAS-mediated extrinsic apoptotic engagement,^13,14^ yet none are rooted directly in genomic markers from CAR-19-treated cases.

In addition to markers of genomic complexity, our WGS results implicated a candidate single-gene alteration, del 3p21.31 containing *RHOA*.^9^ *RHOA* is frequently altered also in newly diagnosed DLBCL, but its roles in pathogenesis remain poorly defined. For example, 3p21.31 deletions occurred in 118/699 (16.9%) of cases in the multi-institution dataset establishing the DLBclass molecular disease clusters,^15^ reflecting consensus across multiple data sets that examined LBCL genomic copy number at resolution.^16–19^ In DLBclass, *RHOA* deletions are a defining feature of cluster 2 (C2) cases, comprising a mix of germinal center B-cell (GCB), activated B-cell (ABC), and unclassified cell of origin (COO).^15^ Non-synonymous *RHOA* mutations also are recurrent, and in DLBclass are a defining feature of C1 (ABC-enriched), occurring in 31/699 (4.4%) overall. These alterations, therefore, implicate loss of function of the Ras homolog family member A (RHOA) protein since only deletions are recurrent, and mutations disrupt RHOA’s interactions with activating guanine-exchange factor (GEF) proteins.^16,17^ Specific functional studies on RHOA in DLBCL, however, are lacking, and its roles are currently defined downstream of other tumor suppressors.^20,21^ Therefore, given poorly defined roles of a gene selected for deletion or deleterious mutations in ∼1 in 5 newly diagnosed DLBCL cases and strong selection for its loss in CAR-T-resistant cases, dedicated studies are highly warranted.

We generated DLBCL RHOA loss-of-function (LoF) systems to assess mechanisms of immune evasion by RHOA-deficient DLBCL, including syngeneic RHOA-deficient lymphoma mouse models. As in prior studies,^22^ complete *RHOA* deletion was not tolerated in DLBCL cells, and our findings therefore characterize it as a haploinsufficient tumor suppressor. These studies represent the first direct functional evaluation of a recurrently deleted LBCL gene linked specifically to poor clinical responses to CAR-19.

## METHODS

### Whole genome sequencing

Raw FASTQ reads were aligned to GRCh38, and somatic single-nucleotide variants (SNVs), short insertion-deletions variants (INDELs), copy-number variants (CNVs) and structural variants (SVs) were called using MGP1000 with additional quality filters.^9^ The dN/dScv method assessed genes under positive selection in r/r. Mutational signatures, single-base substitutions (SBSs), and complex events were defined as described.^9^

### Single-cell RNA-sequencing

A20 tumors were dissociated for droplet-based scRNA-seq on 10x Chromium. 3.5×10^6^ unique barcodes indexed cells with library sequencing by Illumina Novaseq X. Data processing employed Cell Ranger (10x Genomics). Seurat v5.1.0 processed matrices for normalization, scaling, and clustering, followed by Libra v.1.7 aggregation of UMI counts. Cell annotation employed SingleR v.2.4.1 by ImmGen with fine labels and manual validation. Differential expression analysis employed “FindMarkers” (Seurat, min.pct= 0.05).

### Mice and cell lines

NOD.Cg-*Prkdc^scid^ Il2rg^tm1Wjl^*/SzJ (NSG, RRID: MGI:3577020) mice were maintained in pathogen-free facilities. Equal numbers of male and female mice were randomized into untreated and treated conditions. BALB/c (RRID: MGI:2161072) were used for A20 syngeneic tumors.

Cell lines were verified by STR fingerprinting and confirmed mycoplasma negative. RIVA, SU-DHL4, and A20 were cultured in RPMI (10% FBS, 1% penicillin/streptomycin (10,000 units/mL), plasmocin prophylactic (100 mg, Invivogen), and 55 uM 2-ME (A20 only)); OCI-Ly1 in IMDM (20% FBS, 1% penicillin/streptomycin, and plasmocin prophylactic). Cell lines were from ATCC, except RIVA from DSMZ.

### IFN**γ** treatments and priming

Cells at 5×10^5^ cells/mL in triplicate were treated with IFNγ (30 ng/mL, MedChemExpress Cat# HY-P7025, Cat# HY-P7071) for 2 h or 48 h. For the IFNγ primed coculture, cells were seeded at 5×10^5^ cells/mL in triplicate and treated with IFNγ or H_2_0 for 48 h. Cells were cocultured with activated BALB/c T cells (isolated via EasySep Mouse T Cell Isolation Kit (StemCell, 19851A)) in quadruplicate. Percent lysis was determined using luminescence-based cytotoxicity.

### CD19 membrane quantitation

BD Quantibrite™ PE Phycoerythrin Fluorescence Quantitation Kits were used per manufacture’s protocol on 10^5^ cells/replicate.

### In vitro CAR-T cytotoxicity assays

Flow cytometry, luminescence, and impedance-based cytotoxicity protocols and additional details are outlined in Supplementary Appendix.

## RESULTS

### RHOA deficiency drives inflammation-linked CAR-19 resistance in LBCL patients

We interrogated whole genome sequencing (WGS) data from 54 patients with relapsed/refractory (r/r) large B-cell lymphoma (LBCL) treated with anti-CD19 CAR T-cell therapy (CAR-19), 49 of whom were included in our previous study.^9^ We also analyzed WGS from 96 newly diagnosed B-cell lymphoma patients from Pan-Cancer Analysis of Whole Genomes (PCAWG).^23^ Of 150 total cases, 86 were diffuse large B-cell lymphomas (DLBCL), 45 follicular lymphoma/transformed follicular lymphoma, 3 transformed chronic lymphocytic leukemia, 13 Burkitt lymphoma, 2 marginal zone B-cell lymphoma, and 1 post-transplant lymphoproliferative disorder (Supplementary Table 1). Focusing genomic drivers previously associated with clinical outcomes after CAR19, alterations in *RHOA* (mutations or deletions), APOBEC and SBS18 mutational signatures, and chromothripsis were significantly enriched in r/r patients treated with CAR19 (p < 0.05; **Fig. 1A**). All retained significance in predicting early relapse after CAR19 (**Fig. S1A**). Del 3p21.31 or *RHOA* mutations occurred in 23 patients (22% in CAR-19 [n=12]; 14.7% in PCAWG [n=11], **Fig. 1A**). Among these, monoallelic deletions were present in 78%, monoallelic mutations in 17.4%, and biallelic inactivation in one sample (PCAWG). Consistent with DLBclass C2, there was no COO enrichment (p=0.713) (**Fig. S1B**). Although some 3p21.31 deletions also included the *FHIT* tumor suppressor (altered in some digestive-tract tumors)^24^, high-resolution analyses revealed *RHOA* was the minimally deleted region and consistently affected in all cases (**Fig. 1B**).

**Figure 1:**
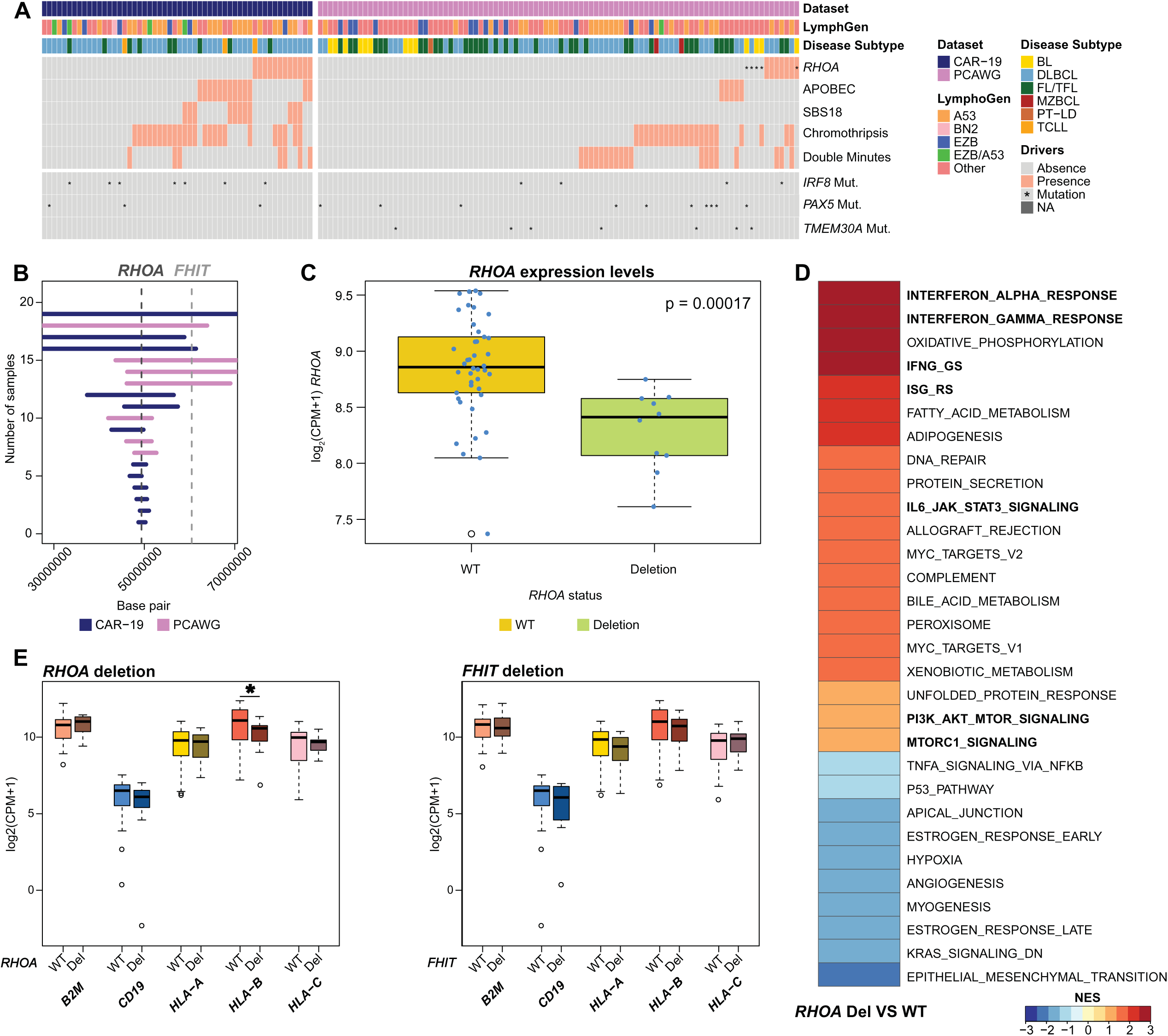
Genomic and transcriptomic analysis in CAR-19 and PCAWG LBCL datasets. **A)** Heatmap displaying genomic drivers associated with poor outcome to anti-CD19 CART cell therapy. Each column represents a patient (CAR-19 and PCAWG datasets), and each row correspond to a genomic driver or annotation. ‘Presence’ and ‘Absence’ are used to identify copy-number variations, mutational signatures, structural variants, and complex events. ‘Mutation’ specifically indicates non-synonymous mutations. The list of driver genes was generated by combining Jain et al. *Blood* 2020 and Sworder et al. *Cancer Cell* 2023. **B)** Segmentation plot highlighting genomic regions involving *RHOA* and *FHIT* deletions across samples from CAR-19 and PCAWG datasets. The y-axis represents the samples, and the x-axis shows the genomic region coordinates. **C)** Boxplot showing the *RHOA* expression levels based on the 3p21.31 deletion status. Each dot represents an individual patient. P-value computed using Wilcoxon test (p<0.05). **D)** Heatmap illustrating significant pathways (FDR<0.05) identified through Gene Set Enrichment Analysis. Pathways of particular interest are shown in bold. **E)** Boxplots comparing expression levels of MHC Class I genes (*HLA-A*, *HLA-B*, *HLA-C*, and *B2M*) and *CD19* in CAR-19 samples with *RHOA* deletions versus WT and *FHIT* deletions versus WT. Significant difference (Wilcoxon test; p=0.035) is marked with an asterisk (*).

We next assessed the consequences of these deletions in bulk transcriptomic data from 52 CAR19-treated LBCL samples with paired WGS. As expected, *RHOA* expression was significantly lower in *RHOA*-deleted compared to WT samples (p=0.00017) (**Fig. 1C**). Additionally, we conducted differential gene expression and gene set enrichment analyses (GSEA). *RHOA*-deleted cases demonstrated multiple LME gene-expression signatures that associate with impaired CAR-19 responses such as JAK/STAT signaling, MYC targets, PI3K-AKT, and mTORC1 signaling (**Fig. 1D**).^7,25,26^ We examined selected immune-related genes, specifically those associated with MHC class I *(HLA-A, HLA-B, HLA-C*, and *B2M*). Interestingly, *RHOA* deletions significantly correlated with reduced expression of *HLA-B* (p = 0.035; **Figure 1E**), a critical component of the MHC class I complex, while *FHIT* deletions had no detectable impact. Overall, we confirm the role del 3p21.31 in predicting LBCL CAR19 outcomes, with *RHOA* as the driver gene involved.

### RHOA-deficient lymphoma cells downregulate CD19 and resist CAR-19 killing

We interrogated the biology of *RHOA* alterations in DLBCL by creating murine and human LoF laboratory systems with *RHOA* shRNA knockdown or CRISPR/Cas9 knockout (RHOA LoF hereafter referring to either sg*RHOA* or sh*RHOA*/sh*Rhoa* systems) (**Fig. 2A**). All were single-cell clones that grew after FACS sorting or serial dilution and were validated via active (GTP-bound) Rho pulldown (**Fig. 2B**). Consistent with previous observations, RHOA LoF DBCL cells showed increased motility in transwell migration assays and aggregation in culture (**Fig. S2A-B**).^21^ sg*RHOA* clones had no proliferative advantage or disadvantage (**Fig. S2C**). Human systems included RIVA, an activated B-cell (ABC) cell of origin (COO) line representing DLBclass C1/LymphGen BN2 subtype (*BCL6* translocation and *NOTCH2* mutation)^18^; OCI-Ly1, a germinal-center B (GCB) COO line; and SU-DHL-4, a CGB COO line derived from C2/A53 subtype.^15^ Murine BALB/c-derived A20 lymphoma was engineered with knockdown of *Rhoa*. Complete loss of *RHOA* never resulted from single-cell selections after sg*RHOA*/Cas9 introduction, consistent with CRISPR screening results reported previously,^22^ implicating *RHOA* as a haploinsufficient tumor suppressor whose complete absence is not tolerated. These systems therefore recapitulate the near exclusive predominance of monoallelic deletions or LoF mutations in patients’ tumors.^23^ Only 2/118 (1.7%) of cases with del 3p21.31 in DLBclass had biallelic deletions, or 0.003% of the 699 cases overall.^15^

**Figure 2:**
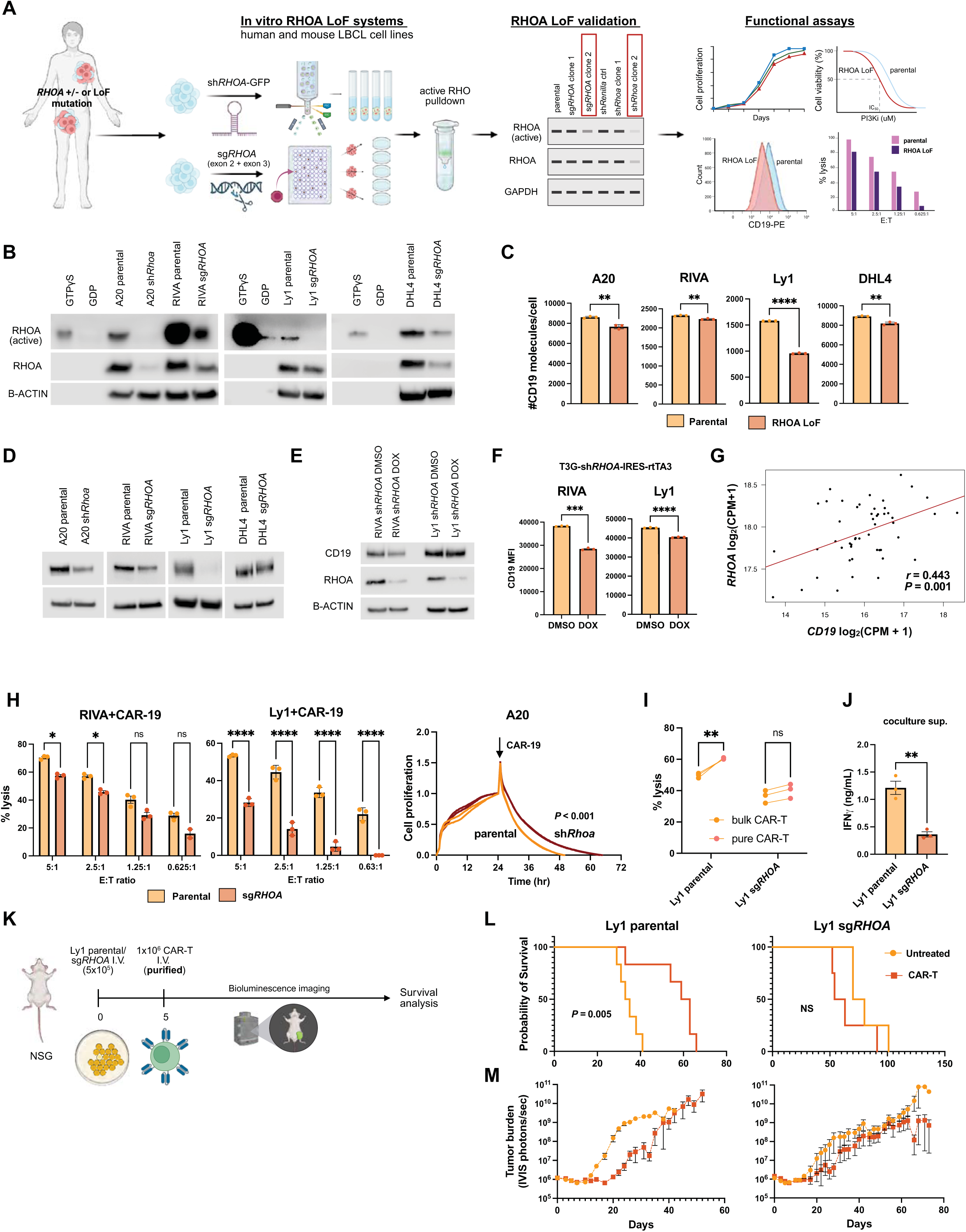
CD19 reduction is responsible for decreased CAR-19 killing of RHOA LoF lymphoma cells in vitro and in vivo. **A)** Schematic of workflow for RHOA LoF cell engineering and validation. **B)** Western blot of human and murine engineered RHOA LoF systems (n=4) revealing decreased active RHOA. GTPyS and GDP serve as positive and negative controls for the pulldown, respectively. Data reflective of two different passages of cells with independent pulldowns. **C)** Phycoerythrin fluorescence quantitation flow cytometry revealing CD19 molecules per cell in RHOA LoF lines. P-value determined via lognormal Welch’s test and reflect mean ± SEM of 3 independent experiments. **D)** Western blot of bulk CD19 protein levels in RHOA LoF cells. Housekeeping genes are sequential as listed and vary from separate experiments. Data reflective of blots from at least two independent passages of cells. **E)** Western blot and **F)** flow cytometry MFI values of RIVA and Ly1 inducible sh*RHOA* knockdown after 10 days of doxycycline (1 µg/mL) exposure. **G)** Correlation between *RHOA* and *CD19* in PCAWG data (n=50 patients, *r* = Pearson’s correlation coefficient). **H)** Left and middle, flow cytometry cytotoxicity assay of RIVA and Ly1 WT and RHOA LoF cells cocultured with purified 4-188 costimulated CD19-CAR-T cells for 6 h. % lysis = 1 -(# targets remaining after coculture/# targets alone). Right, xCELLigence impedance assay of A20 parental or sh*Rhoa* cells with bulk transduced m1928z CAR-T cells (E:T=3:1, CAR-Ts added at 24h, p-value via 2-way ANOVA at a=0.05, data reflects 2 independent experiments). **I)** Luminescence-based cytotoxicity assay of Ly1 WT and RHOA LoF cells cocultured with bulk transduced or purified CD28 costimulated CD19 CAR-T cells for 18 h at 2:1 E:T. P-value derived via 2-way ANOVA at a=0.05. Each connected node reflects an independent experiment. **J)** ELISA of IFNy in supernatants from the Ly1 cocultures with purified CAR-T cells in **I**. P-value determined via Welch’s t test and reflect mean ± SEM of 3 independent experiments. **K**) Schematic of in vivo CAR-T efficacy with Ly1 RHOA LoF system. **L)** Survival curve and **M)** tumor burden measurements of Ly1 parental untreated (n=6), Ly1 parental 1M CAR-T treated (n=6), Ly1 sg*RHOA* untreated (n=4), Ly1

Complete loss of CD19 is uncommon in CAR-19 resistant LBCL, but reduced cell-surface expression may correlates with impaired responses or be found in post-relapse samples.^27–29^ Because genomic drivers of this phenomenon are unknown since *CD19* alterations are rarely detected, we assessed if RHOA LoF could be a driver. By quantitative flow cytometry, we found consistent but variable decline in the number of CD19 surface molecules per cell in RHOA LoF lines (**Fig. 2C**). Western blotting and qPCR showed CD19 downregulation occurred at the level of mRNA expression rather than impaired translation or membrane trafficking (**Fig. 2D, S3A**). Inducible sh*RHOA* cells also demonstrated decreased CD19 after doxycycline induction, while RHOA rescue partially restored expression (**Fig. 2E, S3B**). Likewise, a RHOA-specific inhibitor caused dose-dependent CD19 loss in parental DLBCL cells (**Fig. S3C**). Further, we found strong correlation between *RHOA* and *CD19* transcripts in newly diagnosed DLBCL clinical samples (PCAWG, n=50) (**Fig. 2G**).

To test if RHOA LoF provides intrinsic resistance to CAR-19, we performed coculture cytotoxicity assays. Using flow cytometry, impedance, and luminescence-based cytotoxicity assays, we observed decreased lysis of human and murine RHOA LoF lymphoma cells across a range effector:target (E:T) ratios (**Fig. 2H**). Interestingly, differential killing was greater when purified (GFP+) CAR-19 cells were used as opposed to bulk transduced cells (40-50% GFP+) (**Fig. 2I**), consistent with decreased CAR antigen-specific engagement. We also observed decreased IFNγ in supernatants compared to parental cells (**Fig. 2J**). Target cells themselves did not produce IFNγ (**Fig. S3D**).

We next compared efficacy of CAR-19 cells in vivo using the OCI-Ly1 LoF system that had the greatest loss of CD19 surface density (**Fig. 2K**). Parental OCI-Ly1 cells engrafted by tail vein injection to NSG mice showed significant response to treatment with 10^6^ purified CD28-costimulated CAR-19 cells, including tumor volume reduction and significantly improved survival compared to untreated (p=0.0053, **Fig. 2L-M)**. Animals engrafted with Ly1 sg*RHOA* by contrast experienced no benefit from treatment under the same conditions (p=0.208). Indeed, CAR-T-treated animals if anything died sooner at similar tumor volumes, suggesting CAR-T toxicity is more prominent than efficacy when deployed against these RHOA-deficient tumors, though the trend in OS difference did not reach statistical significance. Collectively, these data demonstrate RHOA deficiency in LBCL cells promotes inherent resistance to killing by CAR-19 cells in vitro and in immunocompromised mice when employing a system with significant decline CD19. The variable degree of CD19 loss across systems, however, left us doubtful this mechanism provides a complete explanation for *RHOA* deletion’s strong association with clinical CAR-19 resistance.

### RHOA deficiency drives activation of and reliance on PI3K-AKT-mTOR signaling

We next assessed cell-intrinsic impact of RHOA LoF in a more unbiased fashion through RNA-sequencing. Analysis of up-regulated gene sets comparing RIVA sg*RHOA* to parental cells revealed PI3K/AKT/mTOR activation, consistent with prior work implicating RHOA in suppressing AKT downstream of S1PR2/P2RY8/Ga13 signaling (**Fig. 3A**).^20^ Immunoblotting confirmed increased AKT activation across systems (**Fig. 3B**). Strikingly, this promoted increased dependence on PI3K/AKT/mTOR signaling, indicated by greater sensitivity to inhibitors of PI3KO (idelalisib), PI3KO/γ (duvelisib), AKT (capivasertib), and mTOR (rapamycin) (**Fig. 3C**), while *RHOA* re-expression rescued idelalisib sensitivity (**Fig. 3D**). Inducible sh*RHOA* RIVA and Ly1 cells also demonstrated significant pAKT abundance after doxycycline exposure (**Fig. 3E**). Drug sensitivity and gene expression data from 18 DLBCL cell lines from the depmap.org database corroborated association between decreased *RHOA* expression and AKT (GSK 690693) and mTOR (A-1065-5) inhibitor sensitivity and a trend that did not reach statistical significance for idelalisib (CAL-101, **Fig. 3F**).^30–31^ Collectively, these data suggest loss of RHOA activity in clinical DLBCL via *RHOA* monoallelic deletions promotes a targetable dependence on the PI3K-AKT-mTOR axis.

**Figure 3:**
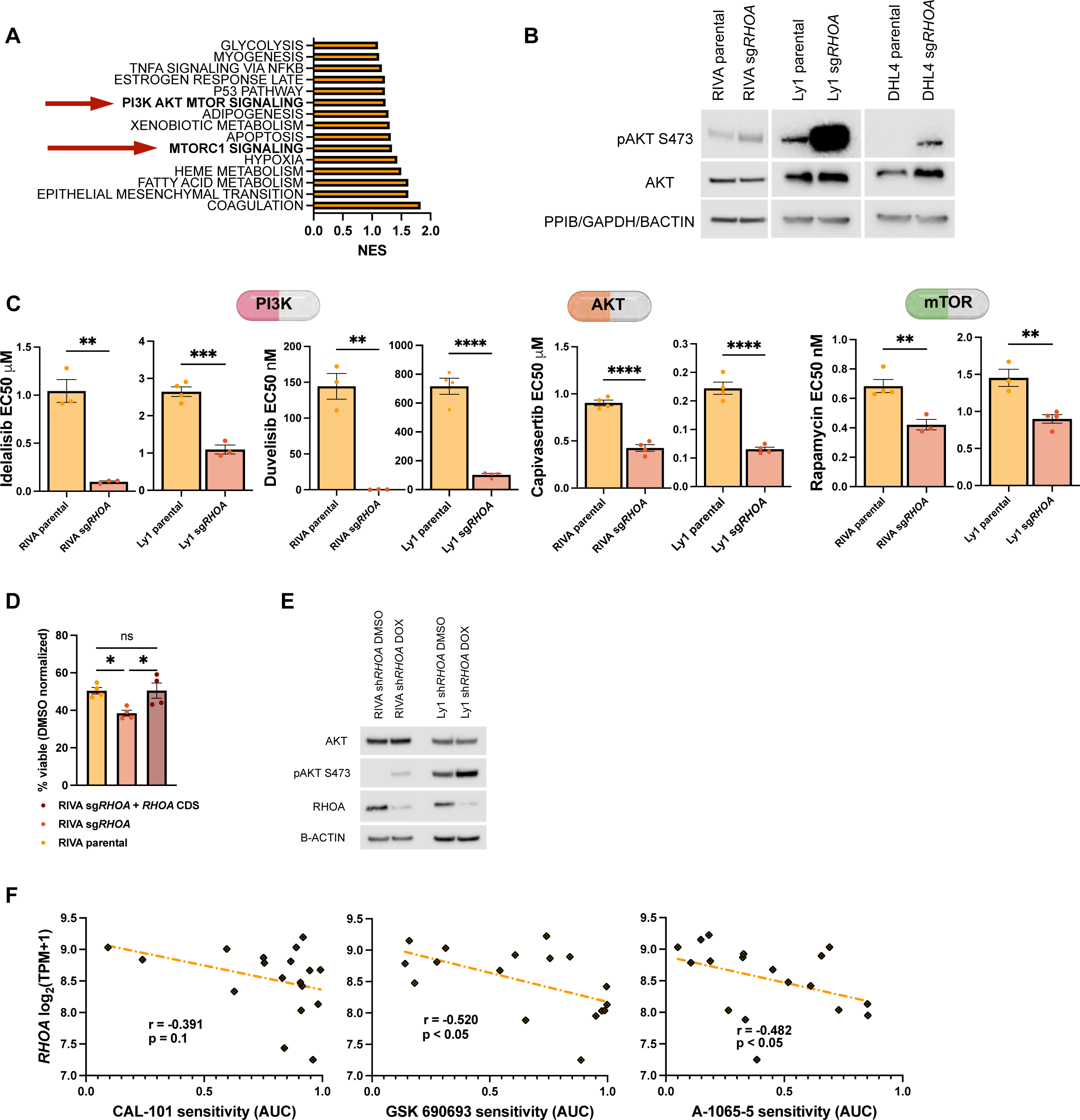
RHOA LoF promotes reliance on PI3K/AKT/mTOR signaling. **A)** GSEA of RIVA RHOA LoF cells compared to RIVA parental cells using the Hallmark gene set (NES>1.0, FDR<0.05). **B)** Western blot of AKT and pAKT levels in LoF cells. Data reflective of blots from at least two independent passages of cells. **C)** EC50 of RHOA LoF cells after treatment with PI3K, AKT, or mTOR inhibitors for 72 hours. Data reflect mean ± SEM of at least 3 independent experiments. **D)** Cell viability (idelalisib 0.315 µM) of RIVA RHOA rescue cells compared to WT and sg*RHOA* cells. Data reflect mean ± SEM of 4 independent experiments. **E)** Western of pAKT induction after 10 days of dox exposure in sh*RHOA* inducible cells. **F)** Drug sensitivity correlations with *RHOA* expression in 18 DLBCL cell lines curated in the DepMap portal. *r* = Spearman’s correlational coefficient. NES - normalized enrichment score, Dox = doxycycline, TPM = transcripts per million. (* p<0.05 **p<0.01 ***p<0.001 ****p<0.0001)

### RHOA loss reshapes lymphoma microenvironments

In contrast to upregulated PI3K signaling, RIVA sg*RHOA* cells exhibited significantly de-enriched interferon gamma and alpha responses (**Fig. 4A-C**). We confirmed specific loss of effector genes *IRF1*, *STAT1*, and *CXCL10* (**Fig. S4A**), and corresponding reduction in CXCL10 secretion (**Fig. S4B**). Activation of chronic IFNγ signaling in LMEs associates with poor CAR-19 clinical responses,^7^ is a driver of the T-exhausted (TEX) LME archetype,^32^ and is significantly enriched in bulk transcriptomic data from 3p.21.31 (*RHOA*) deleted tumors (**Fig. 1D**). Given this, we hypothesized RHOA deficient lymphoma cells blunt tumor cell-intrinsic IFNγ responses to survive chronically inflamed LMEs. To test this, we employed a syngeneic model of RHOA-deficient lymphoma by engrafting parental and sh*Rhoa* A20 cells into strain-matched recipients and harvesting engrafted tumors for single-cell (sc) RNA-seq (**Fig. 4D**). Lymphoma B cells clustered with significant differences between WT and sh*Rhoa* (**Fig. 4E-G**). GSEA revealed mTOR pathway activation and impaired IFNγ response, recapitulating the results for RIVA sg*RHOA* (**Fig. 4H, 3A, 4A**). Gene ontology of biological processes (GO BP) in sh*Rhoa* tumors also revealed significant enrichment of lymphocyte migration and locomotion, consistent with transwell chemotaxis assays (**Fig. S4C, S2A**), and they maintained reduced expression of *Rhoa* and *Cd19* (**Fig. 4I**). Moreover, consistent with suppressed IFNγ responses, we found a striking reduction in antigen processing machinery (*Tap1*) and MHC-I molecules including *B2m* and the HLA-A ortholog *H2-K1* (**Fig. 4J**).

**Figure 4:**
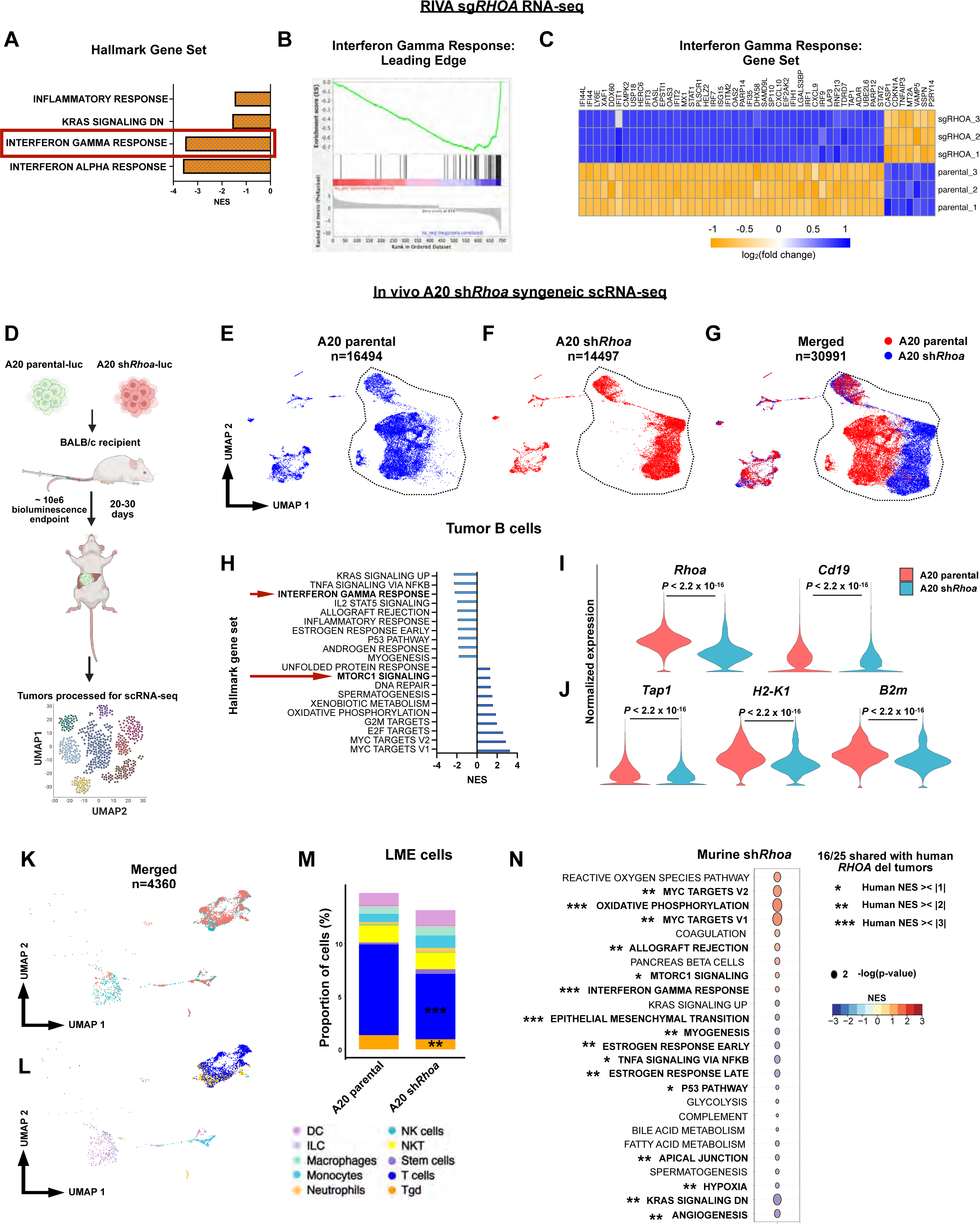
RHOA loss reprograms the lymphoma microenvironment. **A)** GSEA of RIVA RHOA LoF cells compared to RIVA parental cells using the Hallmark gene set (NES<-1.0, FDR<0.05) and corresponding **B)** leading edge analysis and **C)** heatmap of genes in Interferon Gamma Response gene set. **D)** Schematic of syngeneic A20 sh*Rhoa* model of lymphoma and workflow for scRNA-seq analysis. **E), F),** and **G)** UMAP plots of parental and sh*Rhoa* tumors (dashed circle outlines tumor population). **H)** GSEA depicting top 10 upregulated and downregulated gene sets in A20 sh*Rhoa* tumors compared to A20 parental tumors (NOM p-value <0.05). **I)** Expression plots of *RHOA* and *Cd19* in tumor B cells as well as J) *Tap1, H2-K1*, and *B2m* interferon gamma response genes. P-value computed via Wilcoxon test after correcting for multiple hypotheses with Benjamini-Hochberg (BH) method. **K)** UMAP plots of LME cell constituents between genotypes and **L)** annotated. **M)** Proportion of LME cell types in parental or sh*Rhoa* tumors. n = 4 samples/genotype, Wilcoxon test **p<0.01 ***p<0.001. **N)** GSEA analysis for Hallmark gene sets of LME cells in sh*Rhoa* tumors and correlating *RHOA* del (3p.21.31) tumors. Asterisks denote homology to human *RHOA* del GSEA results in Fig. 1D with *NES><|1| **NES><|2| ***NES><|3| pertaining to human data.

We next examined non-tumor LME constituents (**Fig. 4K, 4L**) and observed sh*Rhoa* tumors were somewhat less infiltrated compared to parental (13.16% vs 14.78%, p<0.001) (**Fig. S4D**), largely due to decreased abundance of T cells (6.17% vs. 8.58% (p<0.001) (**Fig. 4M**). To further investigate RHOA-deficient LMEs, we conducted GSEA analysis of pooled LME cells and found remarkable concordance with deregulated gene sets in human *RHOA*-deleted (3p.21.31) tumors (16/25 gene sets with shared deregulation, FDR<0.05, NES>|2|, **Fig. 4N**, compare to **Fig. 1D**). Notably, upregulated oxidative phosphorylation and type 2 interferon responses were apparent, supported also by gene ontology analysis (**Fig. S4E**). *Rhoa* deficiency alone is therefore sufficient to promote microenvironmental features associated with CAR-19 clinical failures.

### Infiltrating T cells in *Rhoa*-deficient tumors demonstrate multiple markers of immune dysfunction

Next, we wanted to assess more specifically T cells found in the microenvironments of *Rhoa*-deficient vs WT tumors. Gene-expression patterns defined nine distinct T-cell clusters (**Fig. 5A-C**, **Fig. S5A-B**). Sh*Rhoa* tumors demonstrated relative depletion of *Cd8a* expression and CD8+ T cells, leading to a CD4:CD8 ratio of nearly twice that of WT (**Fig. 5D-E**). Sh*Rhoa*-enriched CD4+ T cells were *Cd4*^+^*Tbx21*^-^*Cxcr3^-^Eomes*^+^*Ifng*^+^*Prf1*^+^*Gzmb*^+^ Th1-like CD4 cytotoxic T lymphocytes (CD4 CTLs)^33^ and immunosuppressive *Cd4^+^Foxp3*^+^*Gata3*^+^ T regulatory cells (Tregs) **(Fig. 5F-G**). Conversely, sh*Rhoa* tumors showed depletion of *Cd8*a^+^*Tcf7*^+^*Sell*(CD62L)^high^*Cd44*^high^ central memory (CM) cells, a pre-activated undifferentiated phenotype associated with positive CAR-T responses.^5,6^ Consistently, GSEA analysis of T cells revealed multiple enriched pathways for lymphocyte activation and differentiation in the sh*Rhoa* tumors (**Fig. 5H**). Heat map of the differentially expressed genes revealed upregulation of multiple immune checkpoint receptors associated with exhausted T cells (*Pdcd1*, *Lag3*, *Tigit*), lymphocyte recruiting chemokines (*Ccl3*, *Ccl4*), and additional genes that directly promote exhausted phenotypes (*Maf*, *Tox2*) (**Fig. 5I**).^34^ While both CD4 and CD8 subsets expressed increased exhaustion makers, CD8^+^ cells exhibited pronounced multi-marker exhaustion including *Pdcd1*, *Lag3*, *Tigit*, and *Tox* (**Fig. 5J, 5K, Fig. S5C**). Additionally, all T-cell subsets in sh*Rhoa* tumors demonstrated marked downregulation of the AP-1 transcription factors *Jun* and *Fos* that counteract T-cell exhaustion (**Fig. S5D-F**).^35^ *Rhoa* deficiency in tumor cells therefore promoted multiple changes in LME T-cell composition and characteristics associated with immune dysfunction and CAR-T failures.

**Figure 5:**
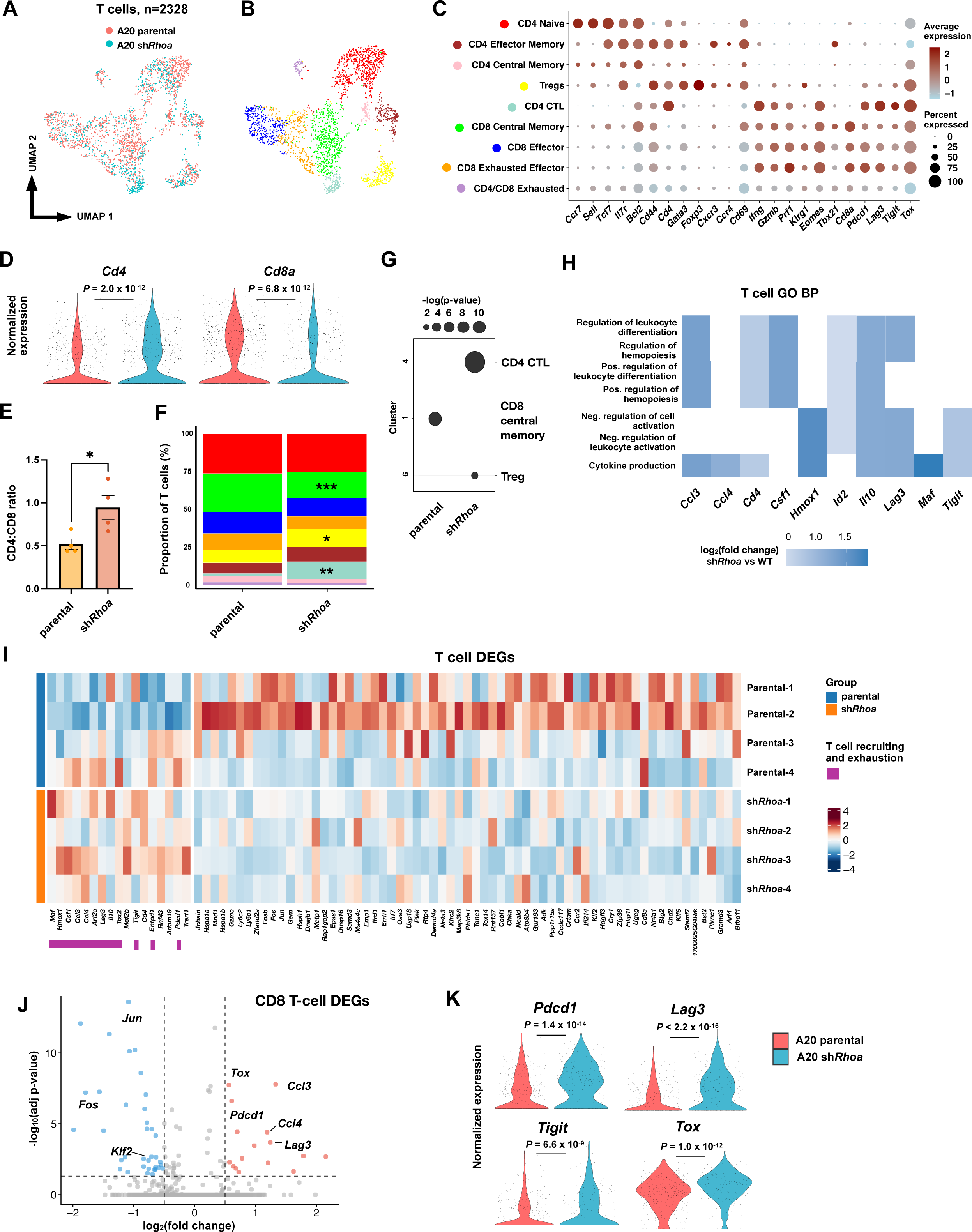
Infiltrating T cells demonstrate CD4 enrichment with multiple markers of exhaustion. **A)** UMAP of bulk annotated T cells across genotypes (n=4 per genotype). **B)** UMAP with annotated T cell subsets described in C). **C)** Expression plot of naïve (*Ccr7*-*Bcl2*), effector (*Cd44*-*Tbx21*), exhausted (*Pdcd1*-*Tox*), and helper/regulatory lineage genes used to define T cell subtypes. Bubble size represents the frequency of each cell type that expresses the indicated gene and color indicates the intensity of expression. **D)** Normalized *Cd4* and *Cd8a* expression across genotypes resulting in an **E)** increased CD4 to CD8 ratio in sh*Rhoa* tumors. P-value for expression plots computed via Wilcoxon test after BH correcting for multiple hypotheses. P-value of ratios derived from unpaired T test, dots represent individual mice (n=4 per genotype) ± SEM. **F)** Proportion of T cell subtypes described in C) (n=4 per genotype). **G)** Dot plot representing abundance/deficiency of certain T cell subsets. Bubble size represents the -log(p-value)) of enrichment/absence. Cluster number outlined in Fig. S5A. **H)** Heatmap of enriched gene sets and corresponding driver genes in pooled sh*Rhoa* T cells. **I)** Heatmap of DEGs in A20 parental and sh*Rhoa* T cells (n=82, adj. p<0.05, FDR<0.05). Heatmap represents Z-scores for each gene. **J)** Volcano plot of DEGs in sh*Rhoa* CD8+ T cells (adj. *P*<0.05, log(FC)>|0.5|). Labeled genes important in T cell recruitment/exhaustion. P-value via Wilcoxon and adjusted with BH method. **K)** Expression plots of known T cell exhaustion genes in sh*Rhoa* CD8 T cells. P-value derived from Wilcoxon test after BH correction.

### CD8 exhausted effectors and CD4 CTLs are central players in dysfunctional sh*Rhoa* LMEs

These findings point to an altered milieu of cell-cell communication in sh*Rhoa* microenvironments promoting immune dysfunction. To assess potential determinants, we further examined individual genes differentially expressed across LME cells. Results reiterated diminished cytotoxic potential in sh*Rhoa* tumors given pronounced decline of genes encoding granzyme production (*Gzmc/d/e/g*) in T-cell subsets, while by contrast there was enhanced expression in various lineages of specific cytokine and chemokine molecules (**Fig. 6A**). Among frequently upregulated genes across LME cell types were *Ccl3*, *Ccl4*, and *Ccl5*, encoding chemokines that attract CCR5+ cells, including cytotoxic T cells and Th1-like CD4 CTLs.^36,37^ In addition, these chemokines promote terminal CD8 T-cell exhaustion in B-cell acute lymphoblastic leukemia.^38^ In both parental and sh*Rhoa* tumors, CD4 CTLs showed the highest *Ccr5* expression(**Fig. 6B**), suggesting mechanism for their increased abundance in sh*Rhoa* lymphomas. We next examined cellular sources of deregulated cytokines and chemokines in sh*Rhoa* LMEs. Global LME expression of *Ccl3* and *Ccl4* was significantly increased in sh*Rhoa* LMEs compared to parental, while the difference for *Ccl5* did not meet statistical significance (**Fig. 6C**). All three showed significantly increased expression in individual cell types, including most notably CD8 exhausted effectors, CD8 CM cells, NKTs, and Tgd cells (**Fig. 6D-G**, **Fig. S6A-C**).

**Figure 6:**
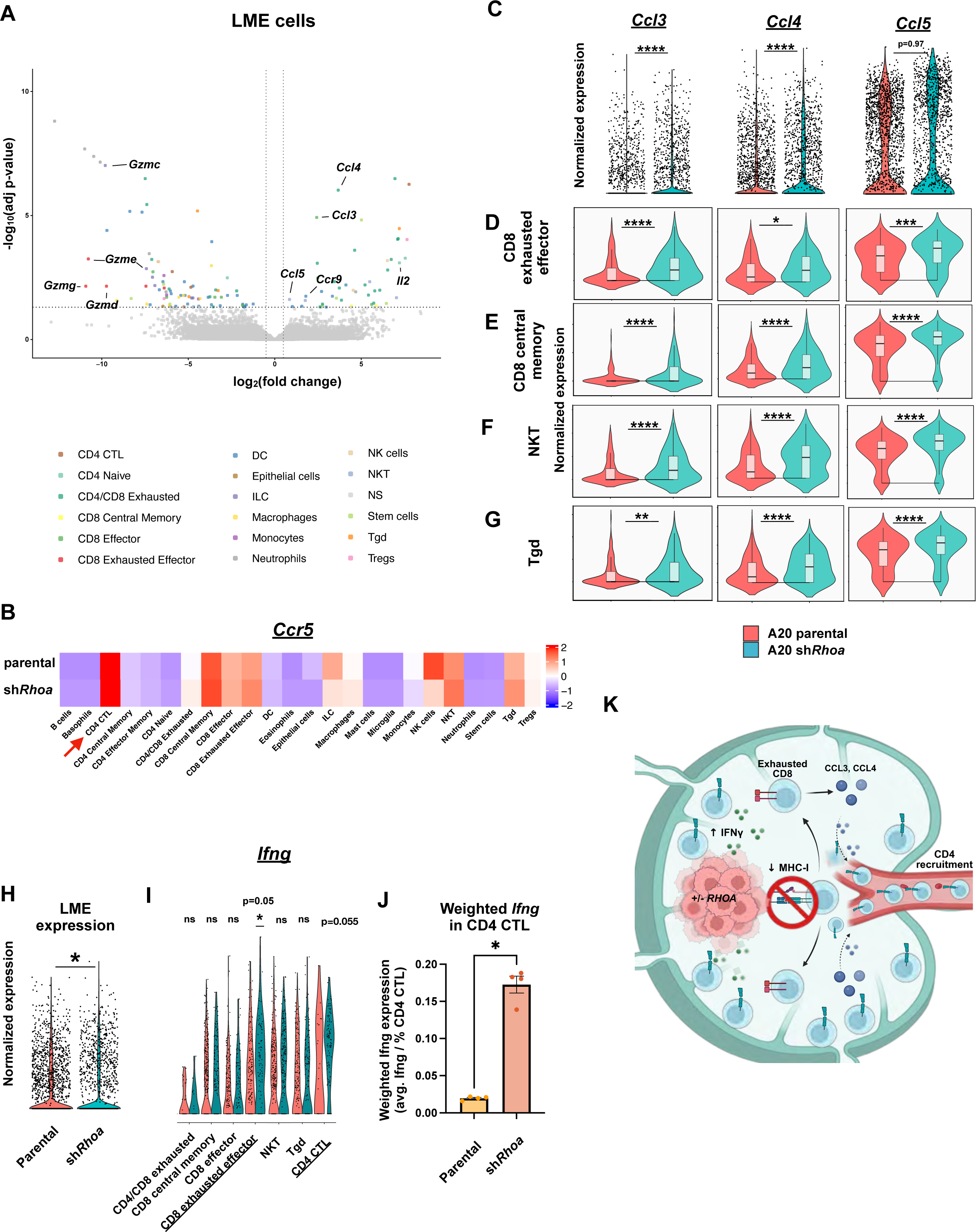
C-C chemokine secretion by exhausted effectors and CD4 CTLs reinforces inflammatory LMEs in shRhoa tumors. **A)** Volcano plot of DEGs in all LME cells (adj. *P*<0.05, log(FC)>|0.5|. Labelled genes important in T cell effector/recruitment. P-value via Wilcoxon and adjusted with BH method. ILC – innate lymphoid cells, DC – dendritic cells, Tgd – gamma delta T cells, NS – not significant. **B)** Heatmap of *Ccr5* expression across all annotated cell types (par – parental, heatmap represents Z-score of *Ccr5* expression). **C)** C-C chemokine expression among all LME cells and **D-G**, selected effector populations in sh*Rhoa* tumors. **H)** Normalized expression of *Ifng* among all LME cells and **I)** selected LME cell types. **J)** Weighted expression plot of total *Ifng* expression in CD4 CTL cells. Weighted expression = (average *Ifng* / % CD4 CTL). % CD4 CTL = 1.8% and 11.5% for parental and sh*Rhoa*, respectively. **K)** Schematic of LME remodeling inflicted by RHOA deficient lymphoma. P-value for all expression plots derived from Wilcoxon test after BH correction * p<0.05 **p<0.01 ***p<0.001 ****p<0.0001.

Simultaneously we found sh*Rhoa* tumor cells significantly under express multiple genes necessary for the cell-surface display of MHC-I (*Tap1*, *H2-K1*, *B2m*, **Fig. 4J**). MHC-I loss by tumors promotes depletion from microenvironments of CD8 CM and other effectors with anti-tumor activities.^39^ Impaired expression of MHC-I machinery in sh*Rhoa* tumor cells occurs despite an enhanced LME IFNγ response, and a global increase in LME *Ifng* expression (**Fig. 6H, Fig. S6D**). CD8 exhausted effectors were the only cell type who’s increased *Ifng* expression was significantly different between sh*Rhoa* and WT LMEs (p<0.05, **Fig. 6I**). CD4 CTLs, however, while not meeting significance on a per-cell basis (p=0.055), were also significantly over-represented in sh*Rhoa* tumors (**Fig. 5F-G**). We therefore determined the compositional weighted mean accounting for both per-cell expression and differential cell prevalence, implicating CD4 CTLs recruited to sh*Rhoa* LMEs as a significant source of increased IFNγ production (**Fig 6J**). Interestingly, tumor B cells themselves did not differentially express any of the specific cytokine or chemokine signaling molecules (**Fig. S6E**), implicating instead their impaired display of MHC-I despite increased presence of IFNγ (validated next) as a key initiating factor in reprogramming LMEs (**Fig. 6K** and discussion).

### Impaired MHC-I display in response to IFN**γ** drives resistance to T cell-mediated killing in RHOA-deficient LBCL cells

We therefore functionally interrogated changes in MHC-I cell surface display by WT and RHOA LoF systems in response to IFNγ. We treated cells with IFNγ (30 ng/mL) and detected MHC-I via the obligate B2M light chain (**Fig. 7A**). Across systems, RHOA LoF promoted impaired MHC-I induction and secretion of IFNγ-induced cytokines CXCL10 and TNFα (**Fig. 7B-D**, **S7A-B**). We examined the dynamics of IFNγ responses by stimulating cells for various timepoints. Western blotting revealed RHOA LoF cell lines have delayed IFNγ responses corresponding to reduced B2M, most pronounced at 12 h (**Fig. 7E-G**). Interferon gamma receptor (IFNGR) levels in RHOA LoF cells were the same or increased compared to parental, showing downstream signaling impairment rather than receptor availability mediates differential responses (**Fig. S7C**). To functionally assess impact on immune-mediated cytotoxicity, we exposed A20 WT and sh*Rhoa* cells to strain-matched T cells with or without 48 hr of priming by IFNγ (**Fig. 7H**). In contrast to a dramatic increase in cell lysis in parental cells after IFNγ stimulation, we observed no lysis in sh*Rhoa* cells with priming (**Fig. 7I**). No difference was observed in proliferation or viability in response to IFNγ (30 ng/mL) between RHOA LoF and parental cells (**Fig. S7D**). RHOA deficiency in DLBCL therefore facilitates avoidance of CD8 immune surveillance by suppressing the tumor cell-intrinsic response to IFNγ, resulting in decreased MHC-I display.

**Figure 7:**
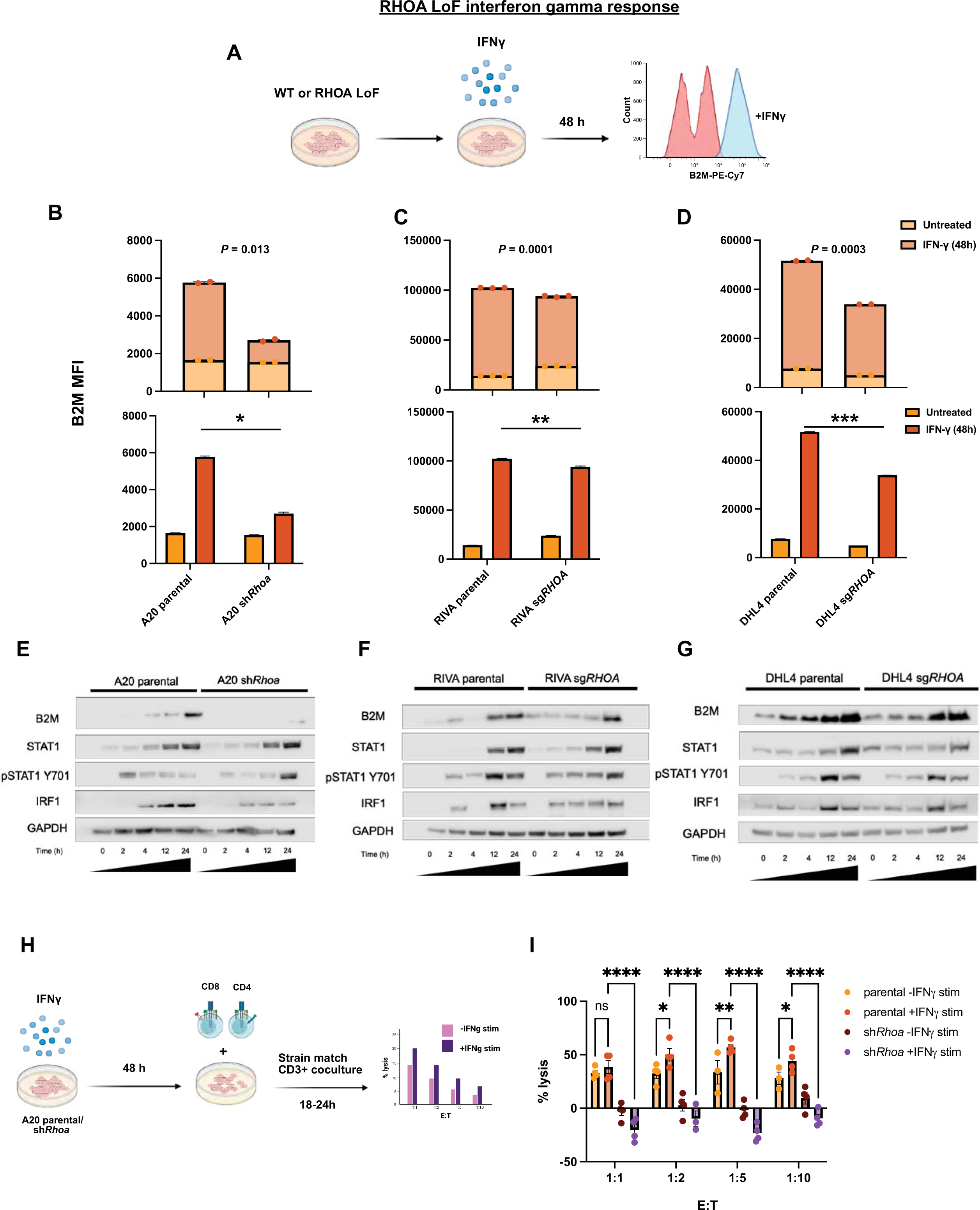
Impaired IFNy responses resulting in deficient MHC-I/B2M induction prohibit T cell-mediated cytotoxicity in RHOA LoF lymphoma. **A)** Schematic of experimental design for assessing B2M induction resulting from IFNy stimulation. **B-D)** Flow cytometry MFI values for parental and RHOA LoF cells after 48 h IFNy stimulation. P-value for both plots derived from log-normal Welch’s T-test. Top plot p-value derived from untreated vs. IFNy treated MFI values. Data reflect mean ± SEM of 2-3 independent experiments **E-G)** Western blot of IFNy time course treatment assessing ISG and B2M protein induction kinetics. Data reflective of blots from at least two independent time course experiments. **H)** Schematic of experimental approach for assessing IFNy response propensity for T cell immunorecognition. **I)** Cytolytic response of A20 parental and sh*Rhoa* cells cocultured with strain-matched T cells for 20 h after IFNy/H20 priming for 48 h. P-value computed via 2-way ANOVA at a =0.05. Data reflect mean ± SEM of 4 independent experiments (* p<0.05 **p<0.01 ***p<0.001 ****p<0.0001).

## DISCUSSION

Current strategies to improve CAR-19 outcomes in LBCL are rooted primarily in CAR-product improvements and largely uninformed by functional studies assessing mediators of CAR19 resistance encoded by lymphoma genomes. Here, we describe for the first time to our knowledge, a single-gene deletion that provides tumor-intrinsic resistance to CAR19. Specifically, reduced activity of the small GTPase, RHOA, both facilitates CD19 antigen escape and promotes formation of immune-dysfunctional microenvironments. *RHOA* has been minimally explored in DLBCL pathogenesis despite alterations occurring in 1 in 5 newly diagnosed patients.^15–19^ Interestingly, previous CRISPR screening implicated *RHOA* as a candidate DLBCL oncogene since its complete deletion was not tolerated.^22^ This is not supported, however, by exclusively LoF genomic alterations. Our data confirm complete RHOA loss is not tolerated: Single-cell selections after sg*RHOA*/Cas9 introduction never yielded viable cells with biallelic deletion (**Fig. 2A-B**). Instead, we define *RHOA* as a haploinsufficient tumor suppressor recurrently selected for reduced gene dosage, reflecting the near-exclusive occurrence of mono-allelic alterations in clinical samples. Most available literature assessing RHOA LoF in DLBCL comes from effects downstream of other tumor suppressors, specifically Ga13 and HGAL,^20,21^ acting to limit cell motility and suppress AKT activation.^41,42^ Increased AKT activation and cell motility were consistent findings due to RHOA LoF in our experiments.

Likely explaining at least in part CAR-19 resistance, we find CD19 antigen decline results from RHOA LoF. While complete loss of CD19 in B-ALL to escape CD19-directed immunotherapies is well documented,^43–45^ the incidence, mechanisms, and implications in DLBCL are less clear. Correlative studies for the ZUMA-7 randomized comparison of axi-cel vs standard chemo/transplant in high-risk second-line patients revealed worse event-free survival in the CAR-T arm for patients with lower pretreatment CD19 by centralized review.^29^ Another series examining more heavily pretreated patients, however, found pretreatment IHC H-scores unpredictive of progression, though some post-relapse samples demonstrated emergence of CD19-low tumors.^46^ We found consistent but variable CD19 downregulation in all RHOA LoF systems generated, one of which had as few as 900 molecules per cell (Ly1 sg*RHOA*, **Fig. 2C**), which unsurprisingly was also most resistant to CAR-19 (**Fig. 2H, K-M**). CD19 downregulation occurred at the level of transcription, and correspondingly there is strong correlation between *RHOA* and *CD19* transcripts in clinical DLBCL cases (**Fig. 2G**). However, the variable impact on CD19 cell surface expression (**Fig. 2C**), carrying variable impact on CAR-19 killing efficacy (**Fig. 2H**), left us skeptical this mechanism provides a complete explanation for *RHOA* deletion’s pronounced association with CAR-19 resistance. In addition, CD19 loss provides poor explanation for the high frequency of *RHOA* deletion in newly diagnosed cases.

We therefore explored the impact of RHOA deficiency in vivo. Syngeneic sh*Rhoa* tumors in immunocompetent hosts replicated to a remarkable degree LME findings in del 3p21.31 LBCL patients (**Fig. 1D, 4N**). In both, IFNγ activation is prominent. Though IFNγ is well described as an activator of antitumor immunity,^47–49^ work in LBCL suggests the cytokine is dispensable for CAR-19 efficacy.^50–52^ Indeed, its effects on LMEs are likely detrimental. A subset of the current authors previously described close associations between clinical CAR-19 treatment failures, tumor interferon signaling, and checkpoint ligand expression on tumor cells.^7^ The current study highlights increased expression of corresponding receptors for these ligands on CD8 T cells in the LME, including PD1, LAG3, and TIGIT. More recently, comprehensive single-cell studies defined three archetypes of LBCL LME, of which one, TEX (T-exhausted), associated with lack of clinical benefit from CAR-19.^32^ IFNγ has prominent roles promoting TEX phenotypes. The fact remains, however, that direct action by IFNγ on tumor cells favors their immune clearance by up regulating MHC-I, to increase visibility to CD8 T cells, and FAS, the cell surface death receptor.^14^ Our results in the context of RHOA deficiency resolve this apparent contradiction. In both human and murine RHOA LoF systems, the response to IFNγ is significantly impaired in malignant cells themselves, manifesting as diminished display of MHC-I in response to IFNγ (**Fig. 7B-D**). We propose this as a core mediator of immune dysfunction in RHOA-deficient LBCLs (**Fig. 6K**). Diminished target recognition by CD8 T cells depletes healthy immune effectors like CD8 CM cells with antitumor potential.^39^ Instead, CD8^+^ T cells in sh*Rhoa* LMEs demonstrate clear stigmata of terminal exhaustion, expressing *Pdcd1, Lag3, Tigit, Tox, Maf*, and others (**Fig. 5K**), markers that consistently correlate with poor outcomes.^5,6^ RHOA-deficient tumors therefore benefit from IFNγ’s detrimental impacts on the overall LME while avoiding the up-regulation of MHC-I that normally would favor their engagement and elimination.

This work provides evidence that RHOA inactivation is a driver of poor CAR-19 efficacy in LBCL by downregulating CD19 and suppressing tumor cell-intrinsic IFNγ responses, limiting CAR-19 direct engagement, protecting from surveilling endogenous CD8^+^ T cells, and fostering dysfunctional LMEs. Our work aligns with a growing body of literature suggesting CAR-19 success is driven as much by an invigoration of endogenous immunity and innate cytotoxic mechanisms as is by CAR-19 direct tumor killing.^8,14,56,62–63^

## Acknowledgements

We wish to thank Dr. Marco Davila for graciously providing the m1928z vector and Mr. Adnan Mookhtiar for sharing expertise of viral transduction techniques. This work was supported by the US Department of Defense Congressionally Directed Medical Research Program (CDMRP) under award HT94252310750 to J.H.S. Research in this publication was performed in part at the Cancer Modeling, Flow Cytometry, and Onco-Genomic Shared Resources (CMSR RRID: SCR022891, FCSR RRID: SCR022501, OGSR RRID: SCR022502) of the Sylvester Comprehensive Cancer Center at the University of Miami, which is supported by the US NIH National Cancer Institute under award P30CA240139. F.L.L. is a Leukemia and Lymphoma Society Scholar in Clinical Research. F.M. is supported by NIH-NCI, LLS and Department of Defense. Sample collection at Moffitt Cancer Center was supported in part by the US NIH National Cancer Institute (P30CA076292). It is also supported by the Mark Foundation, Bankhead-Coley Cancer Research Program. The content is solely the responsibility of the authors and does not necessarily represent official views of the NIH.

## Author Contributions

A.D.N., B.Z., J.Y.S., C.A., J.P.A., M.D.J., F.M., and J.H.S. conceived and designed the project. A.D.N., B.Z., V.V.G.W., C.A.C., Y.E.G., I.S., D.E.T., and P.M. designed or conceptualized experiments. M.V.R., D.G.C., D.B., F.M., and J.H.S. supervised experiments. B.Z. and A.A.Y conducted WGS, RNA-seq, and bioinformatic analysis of patient data. V.V.G.S. manufactured human and mouse CAR-T cells. A.D.N. performed most of the wet lab experiments, as well as the some of the transcriptomic analyses. V.V.G.S., C.A.C., Y.E.G., D.B.C., D.M.S., and A.K.S. performed partial experiments and/or data analysis. N.C.G., N.K., E.R.R., and D.B. performed the mouse experiments. M.D.J., C.A., and F.L.L collected and provided patient data. A.D.N. drafted the manuscript and J.H.S. edited the manuscript with input from all authors, and all authors have approved the final version of the manuscript.

## REFERENCES

1. Neelapu SS, Jacobson CA, Ghobadi A, et al. Five-year follow-up of ZUMA-1 supports the curative potential of axicabtagene ciloleucel in refractory large B-cell lymphoma. Blood. 2023;141(19):2307–2315. doi:10.1182/blood.2022018893

2. Bachy E, Le Gouill S, Di Blasi R, et al. A real-world comparison of tisagenlecleucel and axicabtagene ciloleucel CAR T cells in relapsed or refractory diffuse large B cell lymphoma. Nat Med. Published online 22 September 2022. doi:10.1038/s41591-022-01969-y

3. Jain MD, Spiegel JY, Nastoupil LJ, et al. Five-Year Follow-Up of Standard-of-Care Axicabtagene Ciloleucel for Large B-Cell Lymphoma: Results From the US Lymphoma CAR T Consortium. JCO. 2024;42(30):3581–3592. doi:10.1200/JCO.23.02786

4. Locke FL, Miklos DB, Jacobson CA, et al. Axicabtagene Ciloleucel as Second-Line Therapy for Large B-Cell Lymphoma. N Engl J Med. 2022;386(7):640–654. doi:10.1056/NEJMoa2116133

5. Deng Q, Han G, Puebla-Osorio N, et al. Characteristics of anti-CD19 CAR T cell infusion products associated with efficacy and toxicity in patients with large B cell lymphomas. Nat Med. 2020;26(12):1878–1887. doi:10.1038/s41591-020-1061-7

6. Scholler N, Perbost R, Locke FL, et al. Tumor immune contexture is a determinant of anti-CD19 CAR T cell efficacy in large B cell lymphoma. Nat Med. 2022;28(9):1872–1882. doi:10.1038/s41591-022-01916-x

7. Jain MD, Zhao H, Wang X, et al. Tumor interferon signaling and suppressive myeloid cells are associated with CAR T-cell failure in large B-cell lymphoma. Blood. 2021;137(19):2621–2633. doi:10.1182/blood.2020007445

8. Chen PH, Lipschitz M, Weirather JL, et al. Activation of CAR and non-CAR T cells within the tumor microenvironment following CAR T cell therapy. JCI Insight. 2020;5(12):e134612. doi:10.1172/jci.insight.134612

9. Jain MD, Ziccheddu B, Coughlin CA, et al. Whole-genome sequencing reveals complex genomic features underlying anti-CD19 CAR T-cell treatment failures in lymphoma. Blood. Published online 27 April 2022:blood.2021015008. doi:10.1182/blood.2021015008

10. Cherng HJJ, Sun R, Sugg B, et al. Risk assessment with low-pass whole-genome sequencing of cell-free DNA before CD19 CAR T-cell therapy for large B-cell lymphoma. Blood. 2022;140(5):504–515. doi:10.1182/blood.2022015601

11. Shouval R, Alarcon Tomas A, Fein JA, et al. Impact of TP53 Genomic Alterations in Large B-Cell Lymphoma Treated With CD19-Chimeric Antigen Receptor T-Cell Therapy. JCO. Published online 3 December 2021:JCO.21.02143. doi:10.1200/JCO.21.02143

12. Sworder BJ, Kurtz DM, Alig SK, et al. Determinants of resistance to engineered T cell therapies targeting CD19 in large B cell lymphomas. Cancer Cell. 2023;41(1):210–225.e5. doi:10.1016/j.ccell.2022.12.005

13. Isshiki Y, Chen X, Teater M, et al. EZH2 inhibition enhances T cell immunotherapies by inducing lymphoma immunogenicity and improving T cell function. Cancer Cell. 2024;0(0). doi:10.1016/j.ccell.2024.11.006

14. Upadhyay R, Boiarsky JA, Pantsulaia G, et al. A Critical Role for Fas-Mediated Off-Target Tumor Killing in T-cell Immunotherapy. Cancer Discovery. 2021;11(3):599–613. doi:10.1158/2159-8290.CD-20-0756

15. Chapuy B, Wood T, Stewart C, et al. DLBclass: A Probabilistic Molecular Classifier to Guide Clinical Investigation and Practice in Diffuse Large B-Cell Lymphoma. Blood. Published online 18 December 2024. doi:10.1182/blood.2024025652

16. Monti S, Chapuy B, Takeyama K, et al. Integrative Analysis Reveals an Outcome-Associated and Targetable Pattern of p53 and Cell Cycle Deregulation in Diffuse Large B Cell Lymphoma. Cancer Cell. 2012;22(3):359–372. doi:10.1016/j.ccr.2012.07.014

17. Chapuy B, Stewart C, Dunford AJ, et al. Molecular subtypes of diffuse large B cell lymphoma are associated with distinct pathogenic mechanisms and outcomes. Nat Med. 2018;24(5):679–690. doi:10.1038/s41591-018-0016-8

18. Wright GW, Huang DW, Phelan JD, et al. A Probabilistic Classification Tool for Genetic Subtypes of Diffuse Large B Cell Lymphoma with Therapeutic Implications. Cancer Cell. 2020;37(4):551–568.e14. doi:10.1016/j.ccell.2020.03.015

19. Walker JS, Wenzl K, Novak JP, et al. Integrated genomics with refined cell-of-origin subtyping distinguishes subtype-specific mechanisms of treatment resistance and relapse in diffuse large B-cell lymphoma. Blood Cancer J. 2025;15(1):120. doi:10.1038/s41408-025-01326-5

20. Muppidi JR, Schmitz R, Green JA, et al. Loss of signalling via Gα13 in germinal centre B-cell-derived lymphoma. Nature. 2014;516(7530):254–258. doi:10.1038/nature13765

21. Jiang X, Lu X, McNamara G, et al. HGAL, a germinal center specific protein, decreases lymphoma cell motility by modulation of the RhoA signaling pathway. Blood. 2010;116(24):5217–5227. doi:10.1182/blood-2010-04-281568

22. Reddy A, Zhang J, Davis NS, et al. Genetic and Functional Drivers of Diffuse Large B Cell Lymphoma. Cell. 2017;171(2):481–494.e15. doi:10.1016/j.cell.2017.09.027

23. Aaltonen LA, Abascal F, Abeshouse A, et al. Pan-cancer analysis of whole genomes. Nature. 2020;578(7793):82–93. doi:10.1038/s41586-020-1969-6

24. Simón-Carrasco L, Pietrini E, López-Contreras AJ. Integrated analysis of FHIT gene alterations in cancer. Cell Cycle. 2024;23(1):92–113. doi:10.1080/15384101.2024.2304509

25. Stahl D, Gödel P, Balke-Want H, et al. CSF1R+ myeloid-monocytic cells drive CAR-T cell resistance in aggressive B cell lymphoma. Cancer Cell. 2025;43(8):1476–1494.e10. doi:10.1016/j.ccell.2025.05.013

26. Li X, Singhal K, Deng Q, et al. Large B cell lymphoma microenvironment archetype profiles. Cancer Cell. 2025;0(0). doi:10.1016/j.ccell.2025.06.002

27. Spiegel JY, Patel S, Muffly L, et al. CAR T cells with dual targeting of CD19 and CD22 in adult patients with recurrent or refractory B cell malignancies: a phase 1 trial. Nat Med. 2021;27(8):1419–1431. doi:10.1038/s41591-021-01436-0

28. Plaks V, Rossi JM, Chou J, et al. CD19 target evasion as a mechanism of relapse in large B-cell lymphoma treated with axicabtagene ciloleucel. Blood. 2021;138(12):1081–1085. doi:10.1182/blood.2021010930

29. Locke FL, Filosto S, Chou J, et al. Impact of tumor microenvironment on efficacy of anti-CD19 CAR T cell therapy or chemotherapy and transplant in large B cell lymphoma. Nat Med. 2024;30(2):507–518. doi:10.1038/s41591-023-02754-1

30. DepMap, Broad (2025). DepMap Public 25Q3. Dataset. Published online 2025. depmap.org

31. Arafeh R, Shibue T, Dempster JM, Hahn WC, Vazquez F. The present and future of the Cancer Dependency Map. Nat Rev Cancer. 2025;25(1):59–73. doi:10.1038/s41568-024-00763-x

32. Li X, Singhal K, Deng Q, et al. Large B cell lymphoma microenvironment archetype profiles. Cancer Cell. 2025;0(0). doi:10.1016/j.ccell.2025.06.002

33. Takeuchi A, Saito T. CD4 CTL, a Cytotoxic Subset of CD4+ T Cells, Their Differentiation and Function. Front Immunol. 2017;8:194. doi:10.3389/fimmu.2017.00194

34. Verdeil G. MAF drives CD8+ T-cell exhaustion. OncoImmunology. 2016;5(2):e1082707. doi:10.1080/2162402X.2015.1082707

35. Lynn RC, Weber EW, Sotillo E, et al. c-Jun overexpression in CAR T cells induces exhaustion resistance. Nature. 2019;576(7786):293–300. doi:10.1038/s41586-019-1805-z

36. Nesbeth Y, Scarlett U, Cubillos-Ruiz J, et al. CCL5-Mediated Endogenous Antitumor Immunity Elicited by Adoptively Transferred Lymphocytes and Dendritic Cell Depletion. Cancer Res. 2009;69(15):6331–6338. doi:10.1158/0008-5472.CAN-08-4329

37. Zaunders JJ, Dyer WB, Wang B, et al. Identification of circulating antigen-specific CD4+ T lymphocytes with a CCR5+, cytotoxic phenotype in an HIV-1 long-term nonprogressor and in CMV infection. Blood. 2004;103(6):2238–2247. doi:10.1182/blood-2003-08-2765

38. Zheng J, Zhang Y, Peng X, et al. High expression of CCL3/CCL4/CCL5/CCR5 promotes exhausted CD8+ T cells terminal differentiation and is associated with poor prognosis in pediatric B-ALL patients. Int J Immunopathol Pharmacol. 2025;39:03946320251346823. doi:10.1177/03946320251346823

39. Dhatchinamoorthy K, Colbert JD, Rock KL. Cancer Immune Evasion Through Loss of MHC Class I Antigen Presentation. Front Immunol. 2021;12:636568. doi:10.3389/fimmu.2021.636568

40. Porazzi P, Nason S, Yang Z, et al. EZH1/EZH2 inhibition enhances adoptive T cell immunotherapy against multiple cancer models. Cancer Cell. 2025;0(0). doi:10.1016/j.ccell.2025.01.013

41. Fujisawa K, Fujita A, Ishizaki T, Saito Y, Narumiya S. Identification of the Rho-binding Domain of p160ROCK, a Rho-associated Coiled-coil Containing Protein Kinase *. Journal of Biological Chemistry. 1996;271(38):23022–23028. doi:10.1074/jbc.271.38.23022

42. Vazquez F, Ramaswamy S, Nakamura N, Sellers WR. Phosphorylation of the PTEN Tail Regulates Protein Stability and Function. Mol Cell Biol. 2000;20(14):5010–5018.

43. Sotillo E, Barrett DM, Black KL, et al. Convergence of Acquired Mutations and Alternative Splicing of CD19 Enables Resistance to CART-19 Immunotherapy. Cancer Discovery. 2015;5(12):1282–1295. doi:10.1158/2159-8290.CD-15-1020

44. Cortés-López M, Schulz L, Enculescu M, et al. High-throughput mutagenesis identifies mutations and RNA-binding proteins controlling CD19 splicing and CART-19 therapy resistance. Nat Commun. 2022;13(1):5570. doi:10.1038/s41467-022-31818-y

45. Domizi P, Sarno J, Jager A, et al. IKAROS levels are associated with antigen escape in CD19-and CD22-targeted therapies for B-cell malignancies. Nat Commun. 2025;16(1):3800. doi:10.1038/s41467-025-58868-2

46. Plaks V, Rossi JM, Chou J, et al. CD19 target evasion as a mechanism of relapse in large B-cell lymphoma treated with axicabtagene ciloleucel. Blood. 2021;138(12):1081–1085. doi:10.1182/blood.2021010930

47. Boulch M, Cazaux M, Loe-Mie Y, et al. A cross-talk between CAR T cell subsets and the tumor microenvironment is essential for sustained cytotoxic activity. Science Immunology. 2021;6(57):eabd4344. doi:10.1126/sciimmunol.abd4344

48. Alizadeh D, Wong RA, Gholamin S, et al. IFNγ Is Critical for CAR T Cell-Mediated Myeloid Activation and Induction of Endogenous Immunity. Cancer Discov. 2021;11(9):2248–2265. doi:10.1158/2159-8290.CD-20-1661

49. Wang W, Green M, Choi JE, et al. CD8+ T cells regulate tumour ferroptosis during cancer immunotherapy. Nature. 2019;569(7755):270–274. doi:10.1038/s41586-019-1170-y

50. Larson RC, Kann MC, Bailey SR, et al. CAR T cell killing requires the IFNγR pathway in solid but not liquid tumours. Nature. 2022;604(7906):563–570. doi:10.1038/s41586-022-04585-5

51. Bailey SR, Takei HN, Escobar G, et al. IFN-γ–resistant CD28 CAR T cells demonstrate increased survival, efficacy, and durability in multiple murine tumor models. Science Translational Medicine. 2025;17(801):eadp8166. doi:10.1126/scitranslmed.adp8166

52. Bailey SR, Vatsa S, Larson RC, et al. Blockade or Deletion of IFNγ Reduces Macrophage Activation without Compromising CAR T-cell Function in Hematologic Malignancies. Blood Cancer Discovery. 2022;3(2):136–153. doi:10.1158/2643-3230.BCD-21-0181

53. Ziccheddu B, Saralamma VV, Jain MD, et al. 69 | Escape from Car-Negative Endogenous Cd8 T Cells Drives Resistance to Cd19-Directed Car-T Therapies in Large B-Cell Lymphomas. Hematological Oncology. 2025;43(S3):e69_70093. doi:10.1002/hon.70093_69

54. Cheloni G, Karagkouni D, Pita-Juarez Y, et al. Durable response to CAR T is associated with elevated activation and clonotypic expansion of the cytotoxic native T cell repertoire. Nat Commun. 2025;16(1):4819. doi:10.1038/s41467-025-59904-x

