## Supplement for "*RHOA* Deletion Downregulates CD19 and Promotes Dysfunctional Immune Microenvironments in CAR-T Resistant B-Cell Lymphoma"

#### Supplementary Appendix

Table, Figures, Captions, and Methods

**Supplemental Table 1**

| Characteristic | All patients (N=150) |
|---|---|
| Sex | 60 (40%) |
| Female | 90 (60%) |
| Male | |
| Disease | |
| Diffuse large B-cell lymphoma | 86 (57.3%) |
| Follicular lymphoma/Transformed follicular lymphoma | 46 (30.7%) |
| Transformed Chronic lymphocytic leukemia | 2 (1.3%) |
| Burkitt Lymphoma | 13 (8.7%) |
| Marginal Zone B-cell lymphoma | 2 (1.3%) |
| Post-transplant lymphoproliferative disorder | 1 (0.7%) |
| Dataset | |
| CAR19 | 54 (36%) |
| PCAWG | 96 (64%) |
| Status | |
| Newly Diagnosed | 96 (64%) |
| Relapsed/Refractory | 54 (36%) |
| Median Age | 63 |

### SUPPL. FIGURE 1

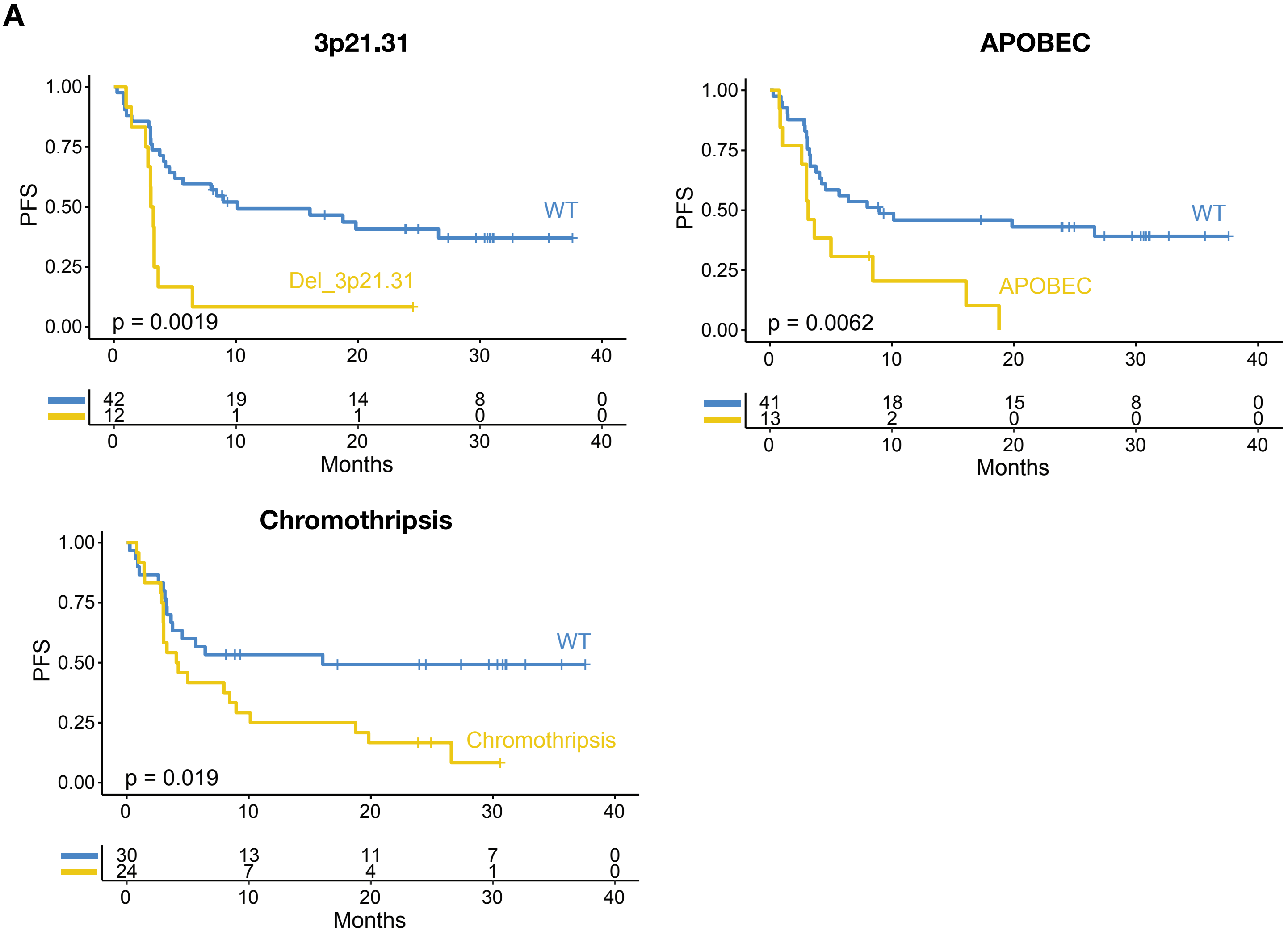


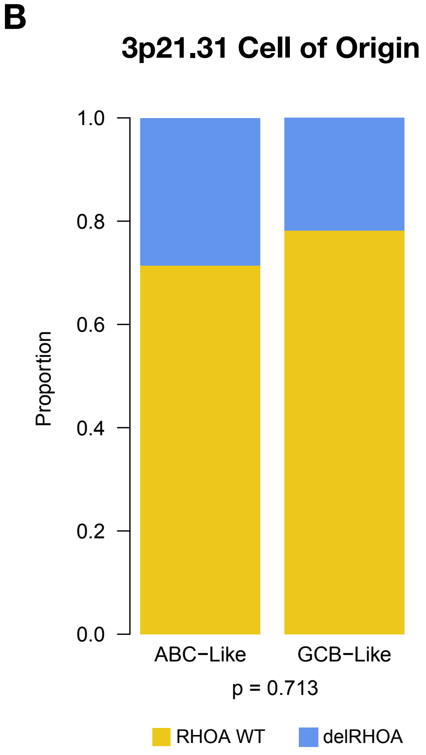

**Supplementary Figure 1: Genomic features associated with CAR19 progression. A)** Kaplan-Meier curve of each independent genomic event implicated in CAR19 progression (p-value determined via Log-rank test with $\alpha = 0.05$). **B)** Cell of origin (COO) classification of CAR19 treated lymphomas with or without *RHOA* (3p.21.31) deletion (n=54). P-value determined with Fisher's exact test at $\alpha = 0.05$.

**SUPPL. FIGURE 2**

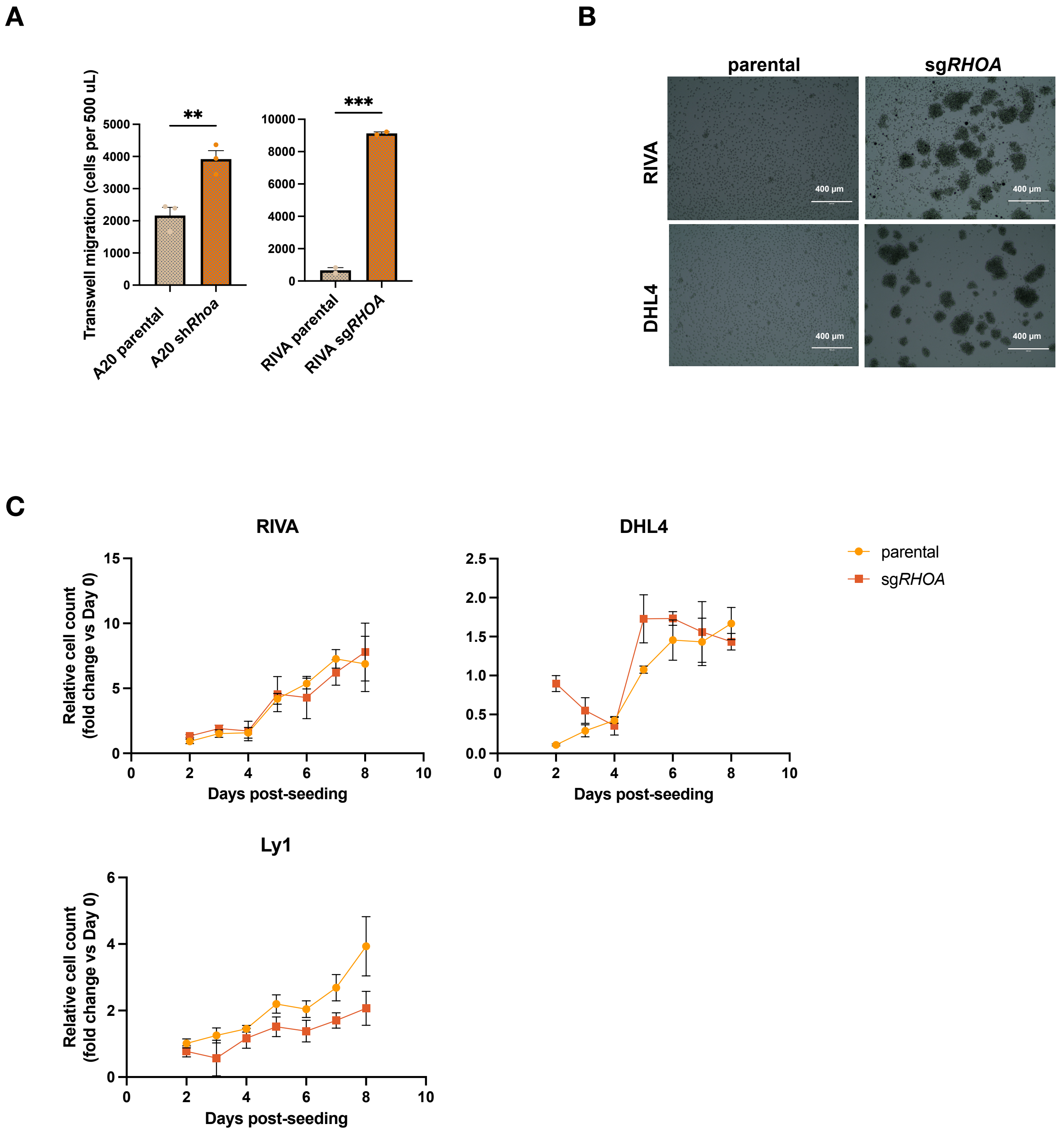

**Supplementary Figure. 2: Phenotypic characterization of RHOA LoF lymphoma cell lines.**

**A)** Cell migration of A20 and RIVA RHOA LoF cells after 4 hr. Total migrated cells were analyzed via flow cytometry by recording the total number of events (cells) in 500 µL of the bottom chamber. Data reflects mean ± SEM 2-3 independent experiments. P-value determined via Welch's T test. **B)** Light microscopy (10x) images of parental and sg*RHOA* cells 48 h post-seeding. **C)** Cell proliferation of parental and sg*RHOA* cells over 8 days. Data reflects mean ± SD of 3 independent experiments.

### SUPPL. FIGURE3

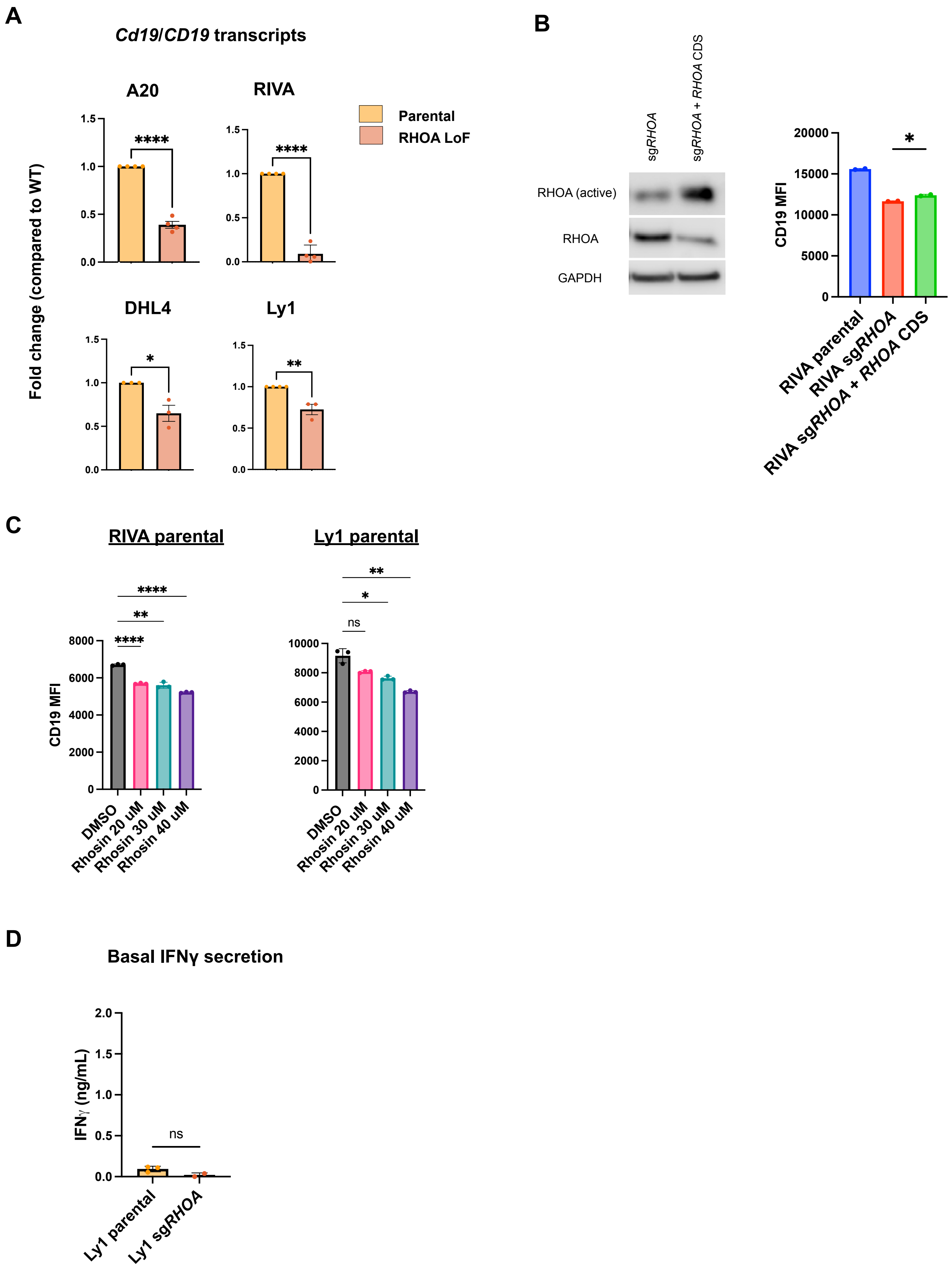

**Supplementary Figure 3: CD19 loss is RHOA-dependent and regulated at the mRNA level.** **A)** qPCR data assessing *CD19* or *Cd19* transcriptional differences between parental and RHOA LoF cells. Fold change determined via delta detla Ct ($2^{-\Delta\Delta Ct}$). P-value determined via Student's t test and represent mean ± SEM of 4 technical replicates, of which is representative of at least 2 independent biological replicates**. B)** Western blot and flow cytometry comparing CD19 abundance in RIVA sg*RHOA* compared to rescue system RIVA sg*RHOA* + *RHOA* CDS. P-value (* p<0.05) determined by lognormal Welch's t test and reflect mean ± SEM of two independent experiments. **C)** Membrane CD19 expression of RIVA and Ly1 parental and RHOA LoF systems after treating with Rhosin for 24 hr. P-value determined via lognormal ordinary one-way ANOVA with α=0.05. Data reflects two independent experiments **D)** IFNγ ELISA of supernatant from wells containing target targets only after 18h. Data reflects 3 independent experiments. (* p<0.05 **p<0.01 ***p<0.001 ****p<0.0001, ns = p>0.05)

### SUPPL. FIGURE 4

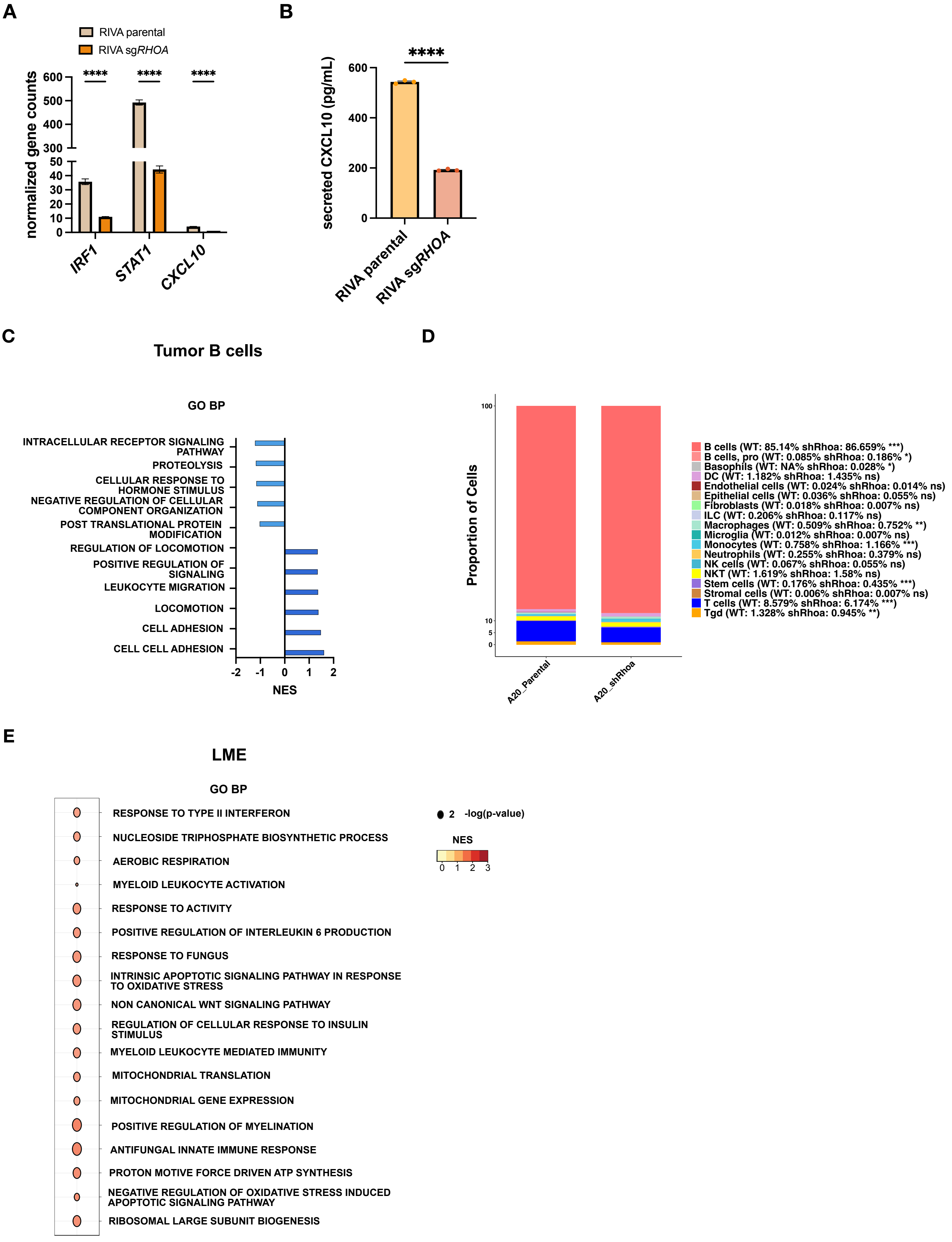

**Supplementary Figure 4: Pathway analysis and LME features of syngeneic sh*Rhoa* tumors. A)** Normalized gene counts of key interferon-stimulated genes from RNA-seq analysis of cultured RIVA sg*RHOA* cells. Counts reflect mean ± SEM of 3 biological replicates **B)** CXCL10 ELISA of supernatants from cultured RIVA parental and sgRHOA cells after 48 h. **C)** GSEA of gene ontology (GO) of biological processes (BP) in B cell tumors of parental and syngeneic sh*Rhoa* mice (n=4 per genotype, adj. P<0.05, FDR<0.05). **D)** Bar plot representing the proportion of each major cell type in A20 parental and sh*Rhoa* syngeneic tumors. P-value determined via Wilcoxon test after correcting for multiple hypotheses with the BH method. Data reflective of n=4 samples/genotype. **E)** GSEA of GO BP in pooled LME cells. Bubble size reflects -log(p-value) and color intensity represents the NES. NES – normalized enrichment score, * p<0.05 **p<0.01 ***p<0.001 ****p<0.0001, ns = p>0.05

### SUPPL. FIGURE 5

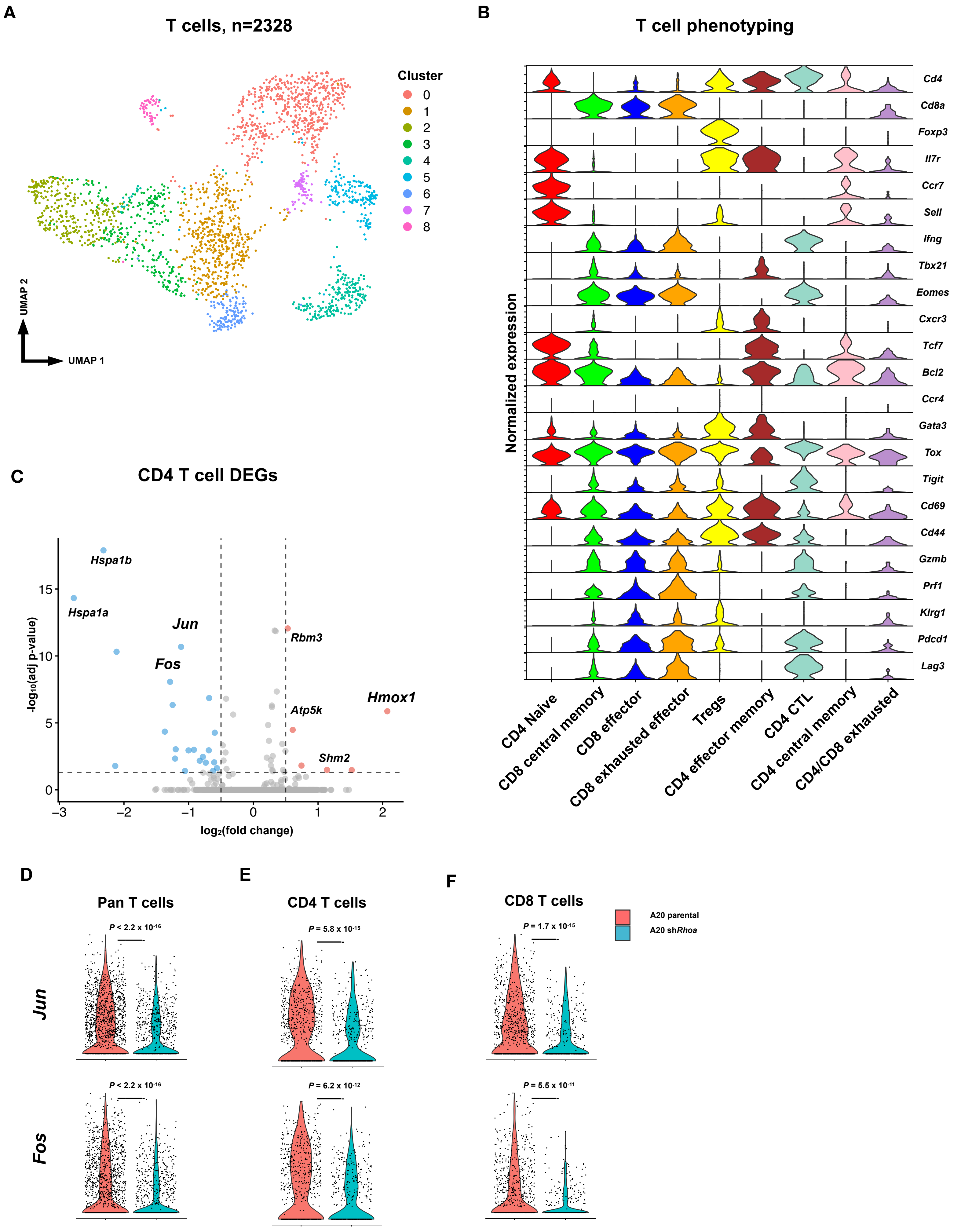

**Supplementary Figure 5: Phenotypic characterization of T cells in scRNA-seq of A20 syngeneic tumors. A)** A20 parental and sh*Rhoa* merged UMAP of bulk annotated T cells with cluster identification (n=4 per genotype). B) Expression violin plots of key genes used define T cell subsets (Tregs – T regulatory cells, CD4 CTL – CD4+ cytotoxic lymphocytes). **C)** Volcano plot of differentially expressed genes (DEGs) in CD4 T cells (adj. $P<0.05$, log(FC)>|0.5|. P-value computed via Wilcoxon and corrected with BH method. Important activation/exhaustion genes are labelled. **D)** Normalized expression of *Fos* and *Jun* in bulk annotated T cells and **E)** CD4 and **F)** CD8 subsets. P-value determined by Wilcoxon and corrected via BH.

### SUPPL. FIGURE 6

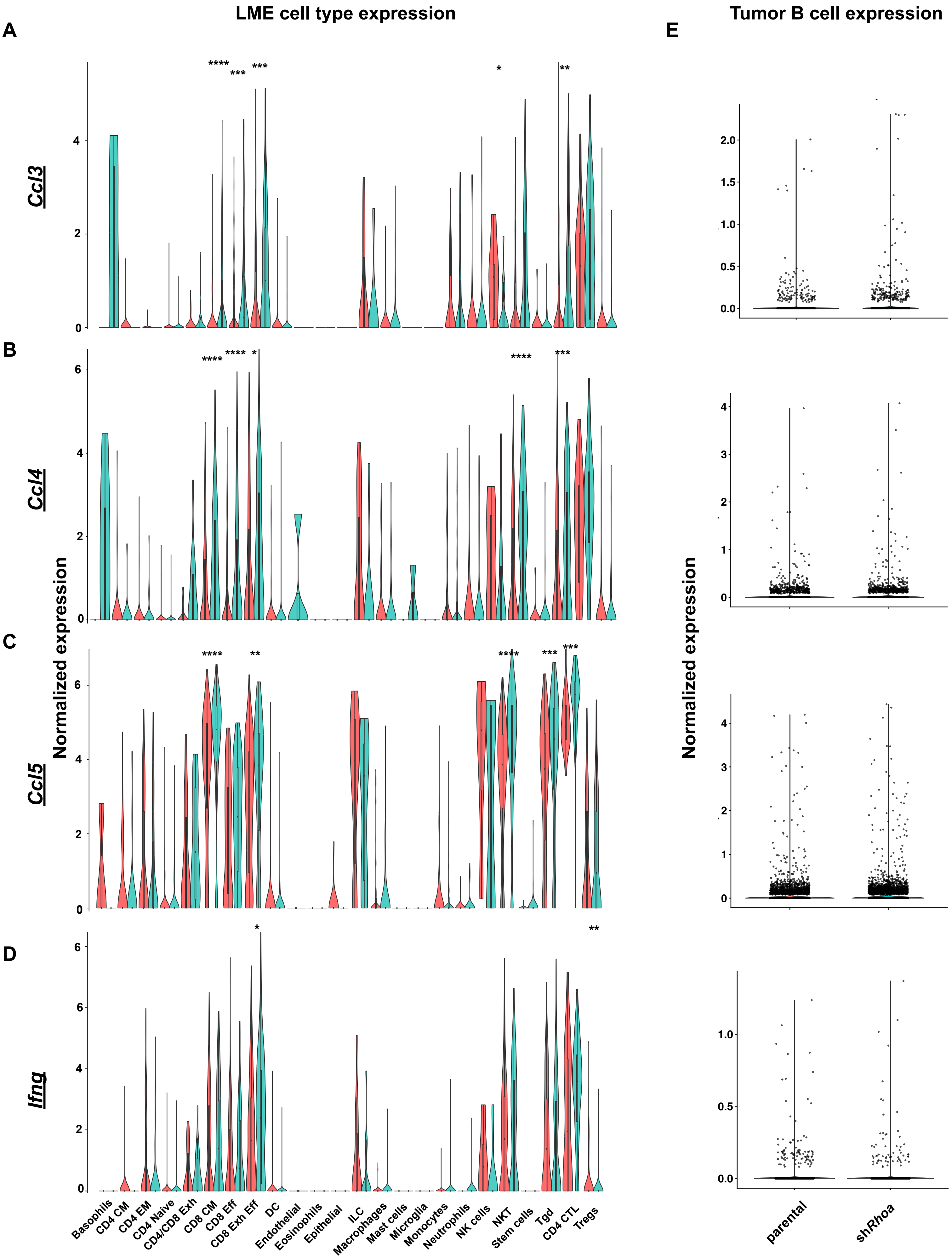

**Supplementary Figure 6: Chemokine expression profiles across LME cells in A20 parental and sh*Rhoa* syngeneic tumors. A)** Normalized expression of *Ccl3*, **B)** *Ccl4*, **C)** *Ccl5*, and **D)** *Ifng* in LME cells. **E)** Normalized expression of respective chemokines in parental and sh*Rhoa* tumor B cells. P-value computed via Wilcoxon and corrected for multiple hypotheses via BH. CM – central memory, EM – effector memory, Exh – exhausted, Eff – effector, DC – dendritic cells, ILC – innate lymphoid cells, Tgd – gamma delta T cells, CTL – cytotoxic lymphocytes. (* $p<0.05$ ** $p<0.01$ *** $p<0.001$ **** $p<0.0001$)

### SUPPL. FIGURE 7

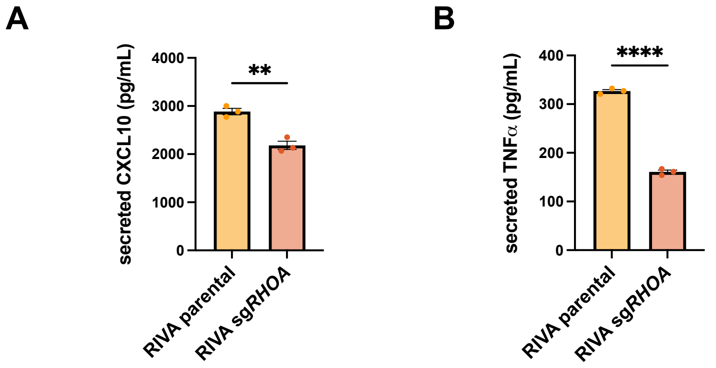


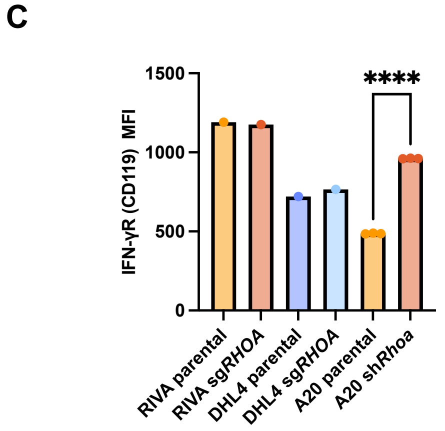


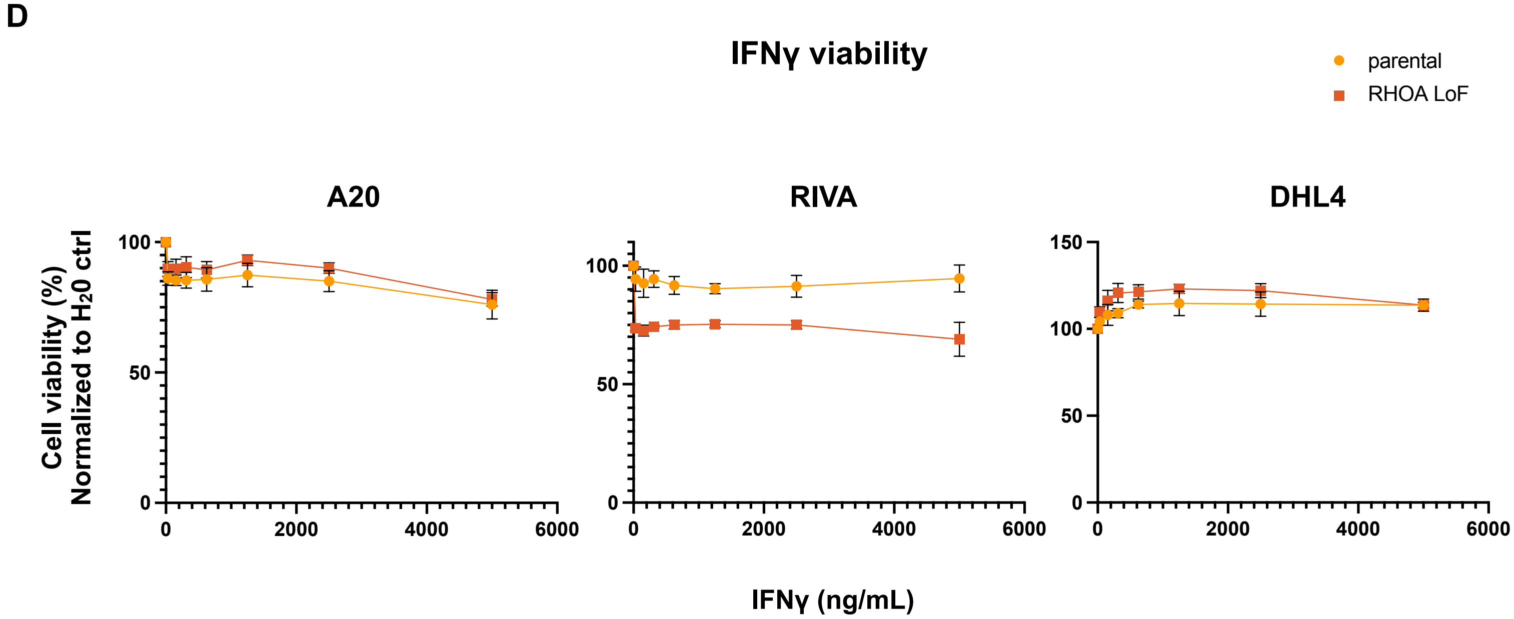

**Supplementary Figure 7: IFNγ-induced cytokine secretion and viability of parental and RHOA LoF lymphoma systems**. **A)** Secretion of proinflammatory CXCL10 and **B)** TNFα by parental and sg*RHOA* cells after 24h IFNγ stimulation (30 ng/mL). Data reflect mean ± SEM of 2-3 independent experiments. P-value determined via Welch's t test. **B)** Interferon gamma receptor (IFNGR, CD119) expression across parental and RHOA LoF systems. P-value determined via lognormal Welch's t test. **C)** Viability of cells cultured for 72 h in serially diluted concentrations of IFNγ. Viability of cells normalized to wells containing cells and $H_20$ control. Data reflects mean ± SD of 4 independent experiments. (**p<0.01 ****p<0.0001)

#### SUPPLEMENTARY METHODS

##### Ethics Statement

The animal experiments were conducted with strict adherence to the institutional guidelines at the University of Miami Miller School of Medicine and follow faithfully the standards of the Institutional Animal Care and Use Committee (IACUC). The Cancer Modeling Shared Resource (CMSR) at the Sylvester Comprehensive Cancer Institute of the University of Miami Miller School of Medicine agree fully to these standards for the entirety of the study involving mice under protocol 21-103-ad06.

##### Mouse models and T cell adoptive transfer

RHOA-deficient model of lymphoma was generated by irradiating 6-12 week BALB/c recipients with 3.5 Gy TBI the morning of lymphoma cell injections. $1x10^6$ A20-luc parental and A20-luc sh*Rhoa* cells were then injected via tail-vein and monitored with IVIS imaging biweekly. Strict endpoints for euthanasia and tumor harvest for scRNA-seq were determined based on loss of 10% body weight, development of hind limb paralysis, or hunched with labored breathing. The in vivo CAR-T efficacy experiment was conducted by engrafting 24 NSG mice 6-12 weeks old with $5x10^5$ luciferase labeled Ly1 parental or Ly1 sg*RHOA* cells. Sample sizes (n=6) were chosen based on prior DLBCL xenografts studies and consistent with previously published CAR-T efficacy preclinical models.[1-2] The primary endpoint was overall survival determined via log-rank test at $\alpha=0.05$ (two-sided). All mice were included in the survival analysis and no mice were excluded or censored for early cytokine release syndrome (CRS) or type IV hypersensitivity CAR-T related deaths. Five days after tumor engraftment, on day 10 of pSLCAR-CD19-28z CAR-T production and two days after sorting GFP+ cells, purified human CAR-Ts were injected at a dose of $1x10^6$ per mouse. Mice were monitored and imaged with IVIS every 2-3 days. Strict endpoints for euthanasia were determined based on loss of 10% body weight, development of hind limb paralysis, bioluminescence imaging (BLI, photons/sec) > $1x10^{11}$, or hunched with labored breathing.

**CAR-T cell generation**

Human – The pSLCAR-CD19-28z 2nd generation CAR construct (Addgene: 135991) was used to generate CD28 costimulated CD19-directed human CAR-T cells. $CD3^+$ T cells were enriched from peripheral blood mononuclear cells (PBMCs) obtained from a single donor (StemCell, #70025.1) using the EasySep™ Human T Cell Enrichment Kit (StemCell, #19051) according to the manufacturer's guidelines. Cells were activated with CD3/CD28 Dynabeads (1:1 cell to bead ratio) (Thermo Fisher, 11131D) and cultured in complete CTS OpTmizer T cell expansion media (Gibco, #A1022101), Complete media: 5% human AB serum, 100 U/mL penicillin/streptomycin, 2mM glutaMAX (Gibco, #A1286001) with 100 IU/mL human IL-2 (R&D, #202-IL-050/CF, 10 ng/mL human IL-7 (StemCell, #78196), and 10 ng/mL human IL-15 (StemCell, #78218). After 48 h, stimulated T-cells were transduced with pSLCAR-CD19-28z lentiviral particles on overnight Retronectin-coated 6-well plates using LentiBOOST (LB) Transduction Enhancer (Revvity, #SB-P-LV-101-10). Briefly, T-cells were mixed with LB and lentiviral particles followed by 10 min incubation at room temperature. Plates were spinoculated at 800 × g for 90 min at 32°C and incubated overnight at 37°C. Fresh complete media was added 24 h post transduction and GFP expression was assessed by flow cytometry 72 h post transduction. CAR-T cells were either sorted or maintained as bulk cultures for downstream experiments and adoptive transfer. Transduction and killing efficacy was confirmed to be similar in at least one other unique PBMC donor. For some assays, a 4-1BB costimulated CD19-directed CAR-T was used (BPS Bioscience, #78171) and cultured in RPMI (10% FBS, 100 IU/mL P/S) and human IL-2 (10 IU/mL, R&D 202-IL-050/CF).

Mouse – For murine CAR-19 generation, spleens from 6–12-week-old BALB/c mice were prepared as single-cell suspensions and lysed with ACK lysis (Thermo Fisher, A1049201) for 4 min. T cells were then isolated with EasySep Mouse T Cell Isolation Kit (StemCell, 19851A), activated with mouse CD3/CD28 Dynabeads (Thermo Fisher, 11452D), and cultured in mRPMI with human IL-2 (80 IU/mL). The next day, previously prepared SFG-m1928z[3] - mCherry retroviral particles were seeded onto a retronectin-coated plate after which T cells were added at $2x10^6$ cells/well and spinfected for 10 min at 1800 RPM at 32 °C. Media was added the following day and bulk transduced m1928z CAR-T cells (~70-80% mCherry+) cells were used in cytotoxic assays.

**RNAseq analysis and Gene Set Enrichment Analysis**

For patient data, raw counts were filtered to remove genes with less than 10 reads in greater than 95% of samples and the trimmed mean of M-values (TMM) normalization was applied. This method estimates a scale factor used to reduce technical bias between samples with different library sizes. For patient samples, cultured RIVA cells, and A20 scRNA we used the DESeq2 (Fig. 1D) or EdgeR (Fig. 3A, 4A) R package to implement the differential expression analysis. P values were corrected for multiple testing using Benjamini-Hochberg false discovery rate method and only genes with FDR<0.05 were considered. Libraries for the cultured RIVA and sg*RHOA* cells were created after 5-10 passages. The Gene Set Enrichment Analysis (GSEA) was performed using the fgsea R package or GSEA v4.3.3 desktop software with the Hallmark and GO BP gene sets collection retrieved from MSigDB database v7.4. The Hallmark gene sets collection was enriched with two INF signatures, ISG.RS and IFNG.GS, previously described to be associated with response to immunotherapy.[4] Genes were ranked using statistics derived from the differential expression analysis, FDR<0.05, p<0.05, and $Log_2FC>|0.5|$.

**sg*RHOA* and sh*Rhoa* LoF Engineered Cell Lines**

CRISPR/Cas9 editing of parental RIVA, Ly1, and DHL4 cells to create RHOA LoF was performed via transient co-nucleofection (Lonza Nucleofector Kit V, VCA-1003) of eSpCas9(1.1)-T2A-GFP-GIntRNA or eSpCas9(1.1)-T2A-mCherry-GIntRNA cloned with sgRNAs targeting exon 2 (TCCGGAAGAAACTGGTGATT) and exon 3 (GATACCGATGTTATACTGAT) of *RHOA* for 24 h followed by FACS sorting of double-positive single cells in 96-well plates. Candidates were expanded and assessed for RHOA LoF via active Rho pulldown assay. Stable knockdown of *Rhoa* in murine A20 lymphoma cells was conducted by designing a 97mer targeting *Rhoa* (TGCTGTTGACAGTGAGCGATGGGTATTCAGTTTTTGAAATAGTGAAGCCACAGATGTATT TCAAAAACTGAATACCCACTGCCTACTGCCTCGGA), PCR amplifying with XhoI and EcoRI restriction sites, and cloning into the mirE-based retroviral shRNA expression vector MLS-E, as previously described.[5] GFP+ cells were sorted, expanded, and validated for *Rhoa* knockdown via active Rho pulldown assay (outlined below).

**Inducible RHOA knockdown and rescue engineered cells**

To generate inducible human *RHOA* knockdown systems a 97mer targeting *RHOA* was designed (TGCTGTTGACAGTGAGCGACTGTAACTACTTTATAACTAATAGTGAAGCCACAGATGTATT AGTTATAAAGTAGTTACAGCTGCCTACTGCCTCGGA), PCR amplified with XhoI and EcoRI restriction sites and cloned into the Tet-ON miR-E (miR-30 variant)-based RNAi vector, LT3GEPIR, as previously described.[5] Lentiviral particles were produced as described below and used to transduce RIVA and Ly1 cells. Cells were selected with puromycin (1μg/mL) and cultured for 2-3 passages (10 days) in doxycycline (1 μg/mL). This duration of induction revealed knockdown of bulk RHOA on Western blot and was used for subsequent experiments. To create RHOA rescue cell lines we engineered the human CRISPR/Cas9 RHOA LoF systems described above via lentiviral introduction of RHOA CDS expression. RHOA ORF GFP-tagged lentiviral particles were purchased (Origene, RC203303L2V) and used to infect cells at MOI >

0.5:1. GFP+ cells were sorted, expanded, and validated for RHOA rescue with active Rho pulldown assay.

**Active Rho pulldown assay and western blotting**

WT and RHOA LoF cells were passaged and seeded at equal densities in 150 mm tissue culture dishes 48 h before creating protein lysates with at least $15 \times 10^6$ cells. At least 250 µg of lysate was subject to treatment with GST-Rhotekin as described by manufacturer (Thermo Scientific, 16116) and subject to GST-Rhotekin pulldown and Western blot analysis. All other western blots were conducted using standard protein extraction protocols and applying an equal amount (~10 µg) of protein per lane. Antibodies used for western blotting include: GAPDH (D16H11) XP® Rabbit mAb (Cell Signaling, Cat# 5174, RRID: AB_10622025), Cyclophilin B Polyclonal Antibody (Thermo Fisher, Cat# PA1-027A, RRID: AB_2169138), RhoA (67B9) Rabbit mAb (Cell Signaling Cat# 2117, AB_10693922), Akt (pan) (11E7) Rabbit mAb (Cell Signaling, Cat# 4685, RRID: AB_2225340), Phospho-Akt (Ser473) (193H12) Rabbit mAb (Cell Signaling, Cat# 4058, RRID: AB_331168), Stat1 Antibody (Cell Signaling, Cat# 9172, RRID: AB_2198300), Phospho-Stat1 (Tyr701) (58D6) Rabbit mAb (Cell Signaling, Cat# 9167, AB_561284), CD19 (Intracellular Domain) (D4V4B) XP® Rabbit mAb (Cell Signaling, Cat# 90176, RRID: AB_2800152), IRF-1 (D5E4) XP® Rabbit mAb (Cell Signaling, Cat# 8478, RRID: AB_10949108), β2-microglobulin (D8P1H) Rabbit mAb (Cell Signaling, Cat#12851, RRID: AB_2716551), β2-microglobulin (F4S6H) Rabbit mAb (Cell Signaling, Cat#86183S).

**Virus generation and transduction**

Retroviruses (m1928z, MLS-E) were generated by transfecting 293T (ATCC, CRL-3216) cells with transfer plasmid and pCL-eco helper plasmid at 3:1 ratios. Lentiviruses (pSLCAR-CD19-28z, LT3GEPIR) were generated by transfecting Lenti-X 293T (Takara, 632180) cells with 4$^{th}$ generation lentiviral transfer plasmids, pMDLg/pRRE (Addgene: 12251), pMD.2G (Addgene:

12259), and pRSV (Addgene: 12253) at a 3:1 ratio of transfer to helper plasmids. Briefly, 293T cells were seeded at $5e^6$ cells in 150 mm dishes. 24-36 h later, DNA was transfected using lipofectamine 3000 (Thermo Fisher L3000001). After 6 h, the transfection media was carefully replaced with fresh DMEM (10% FBS, 100 IU/mL P/S). Supernatant was harvested 48 h later and concentrated with Retro-X or Lenti-X Concentrator (Takara, 631456, 631232) into 100 µL aliquots and frozen in -80 °C for future studies. For most transductions, virus was co-localized onto non tissue culture-treated 6-well retronectin-coated (20 µg/mL, Takara) and 0.5% PBS/BSA blocked plates at 2000xg for 2 h at 32 °C.

**Flow cytometry**

Standard flow cytometry of analysis of isogenic cells was conducted by washing cells ($1e^5$-$5e^5$ cells/sample) with 1mL FACS buffer (Thermo, 00-4222-26) before incubating with Fc receptor blocking cocktails (human, Thermo Fisher 14-9161-73 (2.5 µL/sample), mouse, BD 553141 (1 µL/sample). After 5-15 min. of blocking, antibodies were added at standard manufacturer concentrations. Cells were washed 2x with 1mL FACS buffer and analyzed on Attune NxT flow cytometer. Antibodies used include: PE Mouse Anti-Human CD19 (BD Biosciences, Cat# 555413), PE-Cy™7 Rat Anti-Mouse CD19 (BD Biosciences, Cat# 561739), PerCP/Cyanine5.5 Anti-Mouse CD3 (BioLegend, Cat# 100218), PE/Cyanine7 Anti-Mouse β2-microglobulin (BioLegend, Cat# 154508) and FITC Anti-Human β2-microglobulin (BioLegend, Cat# 316304).

**Flow cytometry cytotoxicity assay**

CAR-T cells were serially diluted in 100 µL of target cell media in triplicates in a 96-well plate. Target cells were labeled with CellTrace Yellow Cell Proliferation Kit (5 µM, Thermo Fisher, C34567), washed, and added to CAR-T containing or control wells at $1x10^4$ cells/100 µL. 6-24 h later plates were analyzed via flow cytometry, with FSC and SSC adjusted on target cells and settings to collect all events in 100 µL. $\% \; lysis = 1 -$

$(\# \; target \; cells \; remaining \; in \; coculture - \# \; cells \; remaining \; in \; target \; only \; wells)$

**Luminescence-based cytotoxicity assay**

Firefly-luciferase labeled Ly1 and Ly1 RHOA LoF cells were cocultured with bulk and purified pSLCAR-CD19-28z CAR-T cells as described in the flow cytometry cytotoxicity assay, without CellTrace labeling. 18-24 h later, plates were centrifuged and wells were washed 1x with PBS and then cocultured cells were lysed with 20 µL 1x lysis buffer (Promega, E1500) for 15 min. Plates were centrifuged and supernatant was transferred 96-well white bottom plates. 100 µL of Luciferase Assay Reagent was added to lysates and plates were immediately read on luminometer. $\% \; lysis = 1 - (luminescence \; of \; coculture - luminscence \; of \; target \; cells \; alone)$

**Impedance-based cytotoxicity assay**

Xcelligence RTCA e-Plates (Agilent, 5469830001) were prepared for A20 lymphoma tethering by coating wells with 0.5 mg/mL anti-B220 overnight at 4 °C. The following day wells were washed, loaded with 50 µL of target cell media analyzed on the RTCA instrument for a brief background reading. Plates were then retrieved from the incubator and A20 cells were seeded at 10,000 cells/well in 100 µL. Instrument was set to collect readings every 30 sec for 24 h, upon which murine CAR-T cells were added at various E:T ratios in 100 µL. Readings continued for 72 h and the raw data (cell index) was plotted and analyzed.

**Transwell migration Assay**

WT and RHOA LoF cells were passaged at equal numbers the day before the experiment and then seeded at $1 \times 10^6$ cells in 300 µL on the top chamber of a 5-micron insert chamber (Sterlitech, 9315012) in serum free RPMI in triplicates in a 24-well plate. The bottom chamber contained serum free RPMI + SDF-1 (CXCL12) (300 ng/mL, R&D Human: 6448-SD-025/CF, Mouse: 460-SD-010/CF) in a total volume of 1 mL. After 4 h, the filter was carefully removed and the bottom chamber containing migrated cells was analyzed via flow cytometry. The total events (cells) collected in 500 μL were recorded, analyzed, and plotted as total cells migrated.

**ELISA**

RIVA basal CXCL10 secretion - cells were seeded at equal densities ($5x10^5$ cells/mL) in triplicates and supernatants (25 μL) were assessed 48 h later using Human IP-10 ELISA Kit Kit (Abcam, ab173194). IFNγ-induced CXCL10 and TNFα (Abcam, ab181421-1) secretion – cells were seeded at $5x10^5$ cells/mL in triplicate and treated with IFNγ for 24 h. Coculture IFNγ assessment – cocultures were performed as described in luminescence-based cytotoxicity assay and supernatant (25 μL) was assessed with Human IFN gamma ELISA Kit (Abcam, ab174443) after 18 h. Quantification was calculated using standard curve and per manufacture's protocol.

**qRT-PCR**

First-strand cDNA synthesis was performed using Oligo (dT) primer and SuperScript III RT (Thermo Fisher, AM5730G, 18080044), according to the manufacturer's protocols. qRT-PCR was performed using a StepOnePlusTM Real-Time PCR System (ThermoFisher Scientific). Relative changes in expression were calculated using the comparative Ct (ΔΔCt) method. 18S rRNA was measured as an internal control for changes in RNA levels. All qPCR reactions were performed using PowerUpTM SYBR® Green Master Mix (Thermo Fisher, #A25742) unless otherwise stated. Primers used include: Human *18S* - F: CAGCCACCCCGAGATTGAGCA R: TAGTAGCGACGGGCGGTGTG, Human *IRF1* - F: ACACAGGCCGATACAAAGCA R: CCATCCACGTTTGTTGGCTG, Human *STAT1* - F: TAATCAGGCTCAGTCGGGGA R: GAAGGTGCGGTCCCATAACA, Human *CD19* - F: TGGGTAATGGAGACGGGTCT R:

TAGCCCTCCCCTTCCTCTTC, Mouse *Rn18S* - F: ATTCGTATTGCGCCGCTAGA R: CCCGGACATCTAAGGGCATC, Mouse *Cd19* - F: TGTCAGCATGCACACATCCT R: GGACCAGGTTTCCCAGGATG, Mouse *Rhoa* F: CGTGGCTGAACTGAGAGTGT R: GCCAACTCTACCTGCTTCCC.

**Drug treatment assay**

Idelalisib, duvelisib, capivasertib, and rapamycin (MedChemExpress) were serially diluted 10 times in 25 µL of growth media with DMSO vehicle in a 384-well plate, in quadruplicates. An equal volume containing 2,500 cells was added and allowed to incubate with drug for 72 h. CellTiter-Glo® reagent (Promega, G7570) was then added to assess viability via luminescence.

**Drug sensitivity correlation with *RHOA* expression**

The DepMap data explorer (depmap.org)[6-7] was accessed and a custom analysis with the Pearson correlation, Sanger GDSC1 data set, *RHOA*, and a curated cell line list was run. The cell lines were obtained by selecting the following disease subtypes: DLBCL- NOS, ACTIVATED B-CELL LIKE, GERMINAL CENTER B-CELL LIKE. Raw data was downloaded and plotted, and correlations analyzed via Pearson's correlation coefficient.

**Pharmacologic RHOA inhibition and CD19 analysis**

250,000 RIVA and Ly1 parental and RHOA LoF cells were seeded in 500 µL in a 24-well plate in triplicates and treated with DMSO or Rho specific inhibitor Rhosin (MedChemExpress, HY-12646) at various concentrations for 24 h. Cells were then subject to flow analysis after Fc block and staining with anti-CD19 (BD, 555413).

**QUANTIFICATION AND STATISTICAL ANALYSIS**

All statistical analyses of transwell migration data, cell viability data, flow cytometry data, qPCR data, drug sensitivity data, ELISA data, cytotoxicity assay data, and mouse survival data were conducted in Prism GraphPad software (v10.6.1, RRID: SCR_002798). WGS, RNA-seq, and scRNA-seq data were analyzed within RStudio (RRID: SCR_000432) using the statistical methods outlined in figure captions.

1. Isshiki Y, Chen X, Teater M, et al. EZH2 inhibition enhances T cell immunotherapies by inducing lymphoma immunogenicity and improving T cell function. *Cancer Cell*. 2024;0(0). doi:10.1016/j.ccell.2024.11.006

2. Upadhyay R, Boiarsky JA, Pantsulaia G, et al. A Critical Role for Fas-Mediated Off-Target Tumor Killing in T-cell Immunotherapy. *Cancer Discovery*. 2021;11(3):599-613. doi:10.1158/2159-8290.CD-20-0756

3. Davila ML, Kloss CC, Gunset G, Sadelain M. CD19 CAR-targeted T cells induce long-term remission and B Cell Aplasia in an immunocompetent mouse model of B cell acute lymphoblastic leukemia. *PLoS One*. 2013;8(4):e61338. doi:10.1371/journal.pone.0061338

4. Jain MD, Zhao H, Wang X, et al. Tumor interferon signaling and suppressive myeloid cells are associated with CAR T-cell failure in large B-cell lymphoma. *Blood*. 2021;137(19):2621-2633. doi:10.1182/blood.2020007445

5. Fellmann C, Hoffmann T, Sridhar V, et al. An optimized microRNA backbone for effective single-copy RNAi. *Cell Rep*. 2013;5(6):1704-1713. doi:10.1016/j.celrep.2013.11.020

6. DepMap, Broad (2025). DepMap Public 25Q3. Dataset. Published online 2025. depmap.org

7. Arafeh R, Shibue T, Dempster JM, Hahn WC, Vazquez F. The present and future of the Cancer Dependency Map. *Nat Rev Cancer*. 2025;25(1):59-73. doi:10.1038/s41568-024-00763-x
